# Single-colony sequencing reveals phylosymbiosis, co-phylogeny, and horizontal gene transfer between the cyanobacterium *Microcystis* and its microbiome

**DOI:** 10.1101/2020.11.17.386995

**Authors:** Olga M. Pérez-Carrascal, Nicolas Tromas, Yves Terrat, Elisa Moreno, Alessandra Giani, Laisa Corrêa Braga Marques, Nathalie Fortin, B. Jesse Shapiro

**Author notes:** O.M.P.C. and N.T. contributed equally to this work. Address correspondence to B. Jesse Shapiro, Olga M. Pérez-Carrascal or Nicolas Tromas, ^1^Département de Sciences Biologiques, Université de Montréal, Campus MIL, 1375 avenue Thérèse-Lavoie-Roux, Montréal, QC, Canada H2V 0B3., or.

## Abstract

Cyanobacteria from the genus *Microcystis* can form large mucilaginous colonies with attached heterotrophic bacteria – their microbiome. However, the nature of the relationship between *Microcystis* and its microbiome remains unclear. Is it a long-term, evolutionarily stable association? Which partners benefit? Here we report the genomic diversity of 109 individual *Microcystis* colonies – including cyanobacteria and associated bacterial genomes – isolated *in situ* and without culture from Lake Champlain, Canada and Pampulha Reservoir, Brazil. We found 14 distinct *Microcystis* genotypes from Canada, of which only two have been previously reported, and four genotypes specific to Brazil. *Microcystis* genetic diversity was much greater between than within colonies, consistent with colony growth by clonal expansion rather than aggregation of *Microcystis* cells. We also identified 72 bacterial species in the microbiome. Each *Microcystis* genotype had a distinct microbiome composition, and more closely-related genotypes had more similar microbiomes. This pattern of phylosymbiosis could be explained by co-phylogeny in two out of the nine most prevalent associated bacterial genera, *Roseomonas* and *Rhodobacter*, suggesting long-term evolutionary associations. *Roseomonas* and *Rhodobacter* genomes encode functions which could complement the metabolic repertoire of *Microcystis*, such as cobalamin and carotenoid biosynthesis, and nitrogen fixation. In contrast, other colony-associated bacteria showed weaker signals of co-phylogeny, but stronger evidence of horizontal gene transfer with *Microcystis*. These observations suggest that acquired genes are more likely to be retained in both partners (*Microcystis* and members of its microbiome) when they are loosely associated, whereas one gene copy is sufficient when the association is physically tight and evolutionarily long-lasting.

## Introduction

Cyanobacteria occur naturally in aquatic ecosystems, often multiplying into harmful blooms and producing a diversity of toxins, which can cause severe human illness^1^. Many cyanobacteria and eukaryotic algae grow in mucilaginous colonies surrounded by a zone, called the phycosphere, rich in cell exudates, where metabolites are exchanged between numerous microorganisms^2,3^. In this microhabitat, the interactions between cyanobacteria and associated bacteria (AB) might include mutualism (with all partners benefitting), competition (with all partners competing for resources), antagonism (inhibiting one of the partners), commensalism (with one partner benefitting) and parasitism (with one partner benefitting at the expense of the other)^3–5^. However, the drivers shaping these associations are largely unknown. In some cases, AB may enhance algal or cyanobacterial growth^6,7^, aiding phosphorus acquisition in *Trichodesmium*^8,9^. Understanding the contributions of AB to cyanobacterial growth and toxin production has implications for our ability to predict and control harmful blooms.

*Microcystis* is a globally-distributed, often toxigenic bloom-forming freshwater cyanobacterium, which forms macroscopic mucilaginous colonies. These colonies offer a nutrient-rich habitat for other bacteria, while also providing physical protection against grazers^10–12^. The *Microcystis* colony microbiome is distinct from the surrounding lake bacterial community, enriched in Proteobacteria and depleted in Actinobacteria^13,14^. The microbiome composition has been associated with temperature, seasonality, biogeography, *Microcystis* morphology and density^13,15–17^. Lab experiments show the potential for AB to influence *Microcystis* growth and colony formation^18–21^. Yet it remains unclear whether such interactions are relevant in natural settings, and if they are the product of long-term associations over evolutionary time.

Phylosymbiosis, a pattern in which microbiome composition mirrors the host phylogeny^22^, provides a useful concept for the study of host-microbiome interactions. Phylosymbiosis could arise from some combination of (1) vertical transmission of the microbiome from parent to offspring, resulting in co-speciation and shared phylogenetic patterns (co-phylogeny), (2) horizontal transmission of the microbiome, but with strong matching between hosts and microbiomes at each generation, and (3) co-evolution, in which hosts and microbiomes mutually impose selective pressures and adapt to each other. Distinguishing the relative importance of these three possibilities can be challenging, but in all cases the associations between hosts and microbiomes are non-random. Phylosymbiosis is typically studied between plant or animal hosts and their microbiomes^23–25^ but *Microcystis* could also be considered a host, since it constructs the mucilage environment – although it is unclear to what extent it selects its AB or *vice versa*. *Microcystis* colonies are more open to the outside environment compared to mammalian guts, for example. Consequently, they might behave more like coral mucus^25^ or other animal surfaces which seem to show weaker phylosymbiosis than guts^26^. The enclosed nature of animal guts reduces dispersal of microbiomes and favours vertical transmission, potentially leading to co-phylogeny without the need to invoke co-evolution^27^. In contrast, metagenomic sequencing suggests *Microcystis* and its microbiome are globally distributed^16^, making it unlikely that phylosymbiosis could arise due to common biogeography of *Microcystis* and its microbiome. On the other hand, *Microcystis* may be geographically structured on shorter evolutionary time scales, due to local adaptation or clonal expansions, and *Microcystis* genotypes might have distinct phenotypic characteristics that could select for distinct microbiomes^28,29^. Phylosymbiosis studies to date are biased toward the gut relative to external host compartments^22^, and *Microcystis* colonies provide an ideal model of a more ‘external microbiome’.

Previous studies of the *Microcystis* microbiome have used either culture-independent metagenomics from lakes, a bulk biomass collection method which cannot resolve fine-scale spatial interaction within colonies (*e.g.*,^16^), or culture-based studies of *Microcystis* isolates, which have found host-microbiome divergence according to phosphorous gradients and taxonomy^30^, but may not be representative of the natural diversity of *Microcystis* or AB as they occur in nature. To combine the strengths of both these approaches, we developed a simple method for isolating individual *Microcystis* colonies directly from lakes, followed by DNA extraction and sequencing without a culture step^29^. Here we applied this method to 109 individual colonies from Lake Champlain, Canada and Pampulha Reservoir, Brazil, yielding 109 *Microcystis* genomes and 391 AB genomes.

Our findings reveal an expanded *Microcystis* genotypic diversity, and a *Microcystis* colony microbiome shaped by the host genotype, resulting in a significant signature of phylosymbiosis. We inferred co-speciation of *Microcystis* with two of the most prevalent genera in its microbiome (*Rhodobacter* and *Roseomonas*) suggesting evolutionarily stable associations. We also inferred extensive horizontal gene transfer (HGT) events among *Microcystis* and its microbiome, mainly involving lower-fidelity partners than *Rhodobacter* and *Roseomonas*. Overall, our results suggest ecologically and evolutionarily stable associations between *Microcystis* and members of its microbiome.

## Results

### Genotypic diversity of *Microcystis* colonies in Lake Champlain and Pampulha Reservoir

To study the relationship between *Microcystis* and its AB in natural settings, we sequenced 109 individual *Microcystis* colonies from 16 lake samples (82 colonies from Lake Champlain, Quebec, Canada and 27 from Pampulha Reservoir, Minas Gerais, Brazil; Supplementary Table 1). *Microcystis* genomes were assembled and binned separately from AB genomes (Methods), which we will describe below. Consistent with our previous study of *Microcystis* isolate genomes^29^, nearly all *Microcystis* genomes share ≥95% average nucleotide identity (ANI), with the exception of 14/53,381 genome pairs with ANI <94.5%. The 95% ANI threshold is typically used to define bacterial species, but we previously found significant phylogenetic substructure above 95% ANI, coherent with multiple species or ecotypes within *Microcystis*^29^. Consistent with such fine genetic structure within our sampled colonies, we identified 18 monophyletic, closely-related genotypes of *Microcystis* (≥99% ANI; Supplementary Table 2 and Fig. 1). These genotypes (highlighted clades in Fig. 1) are nested within the phylogeny of 122 isolate genomes previously sampled from North America, Brazil, and worldwide. However, only two genotypes (G05 and G10) have been observed in culture previously, possibly due to the fine-grained definition of genotypes (≥99% ANI) combined with undersampling of natural diversity in culture collections^31^. Consistent with previously observed biogeographic patterns between North and South America^29^, we found 14 genotypes unique to Lake Champlain, and four unique to Pampulha, with no genotypes found in both locations.

**Table 1.** Co-phylogeny analysis between *Microcystis* and the nine most prevalent associated bacterial genera within the *Microcystis* microbiome.

| Associated bacteria (AB) genus | Number of species per genus | Number of AB genomes used in the phylogeny | Prevalence of AB in colonies from Canada and Brazil | Correlation with *Microcystis* in Canada metagenomes ($r^2$) | ParaFit test ($P$-values) |
| --- | --- | --- | --- | --- | --- |
| *Phenylobacterium* | 5 | 60 | 73.40% | 0.759 * | 0.072 (0.008) |
| *Roseomonas* | 13 | 36 | 70.64% | 0.835 * | 0.009** (0.001) |
| *Rhodobacter* | 4 | 34 | 46.79% | 0.779 * | 0.0018** (0.0002) |
| *Methylobacterium* | 3 | 29 | 44.04% | 0.809 * | 0.729 (0.081) |
| *Pseudanabaena* | 2 | 20 | 43.12% | 0.766 * | 0.153 (0.017) |
| *Rhodocyclaceae bacterium* G1 | 2 | 19 | 39.45% | 0.769 * | 0.225 (0.025) |
| *Rhodocyclaceae bacterium* G2 | 2 | 21 | 31.19% | 0.776 * | 5.355 (0.595) |
| *Chitinophagaceae bacterium* | 3 | 22 | 26.60% | 0.795 * | 0.081 (0.009) |
| *Cytophagales bacterium* | 3 | 16 | 22.94% | 0.740 * | 0.702 (0.078) |

\* significant correlation coefficients ( $Q < 0.01$ ).
\*\* significant $P$ -values ( $P < 0.01$ ) (Bonferroni correction). Uncorrected $P$ -values are shown between parentheses.

**Figure 1.**
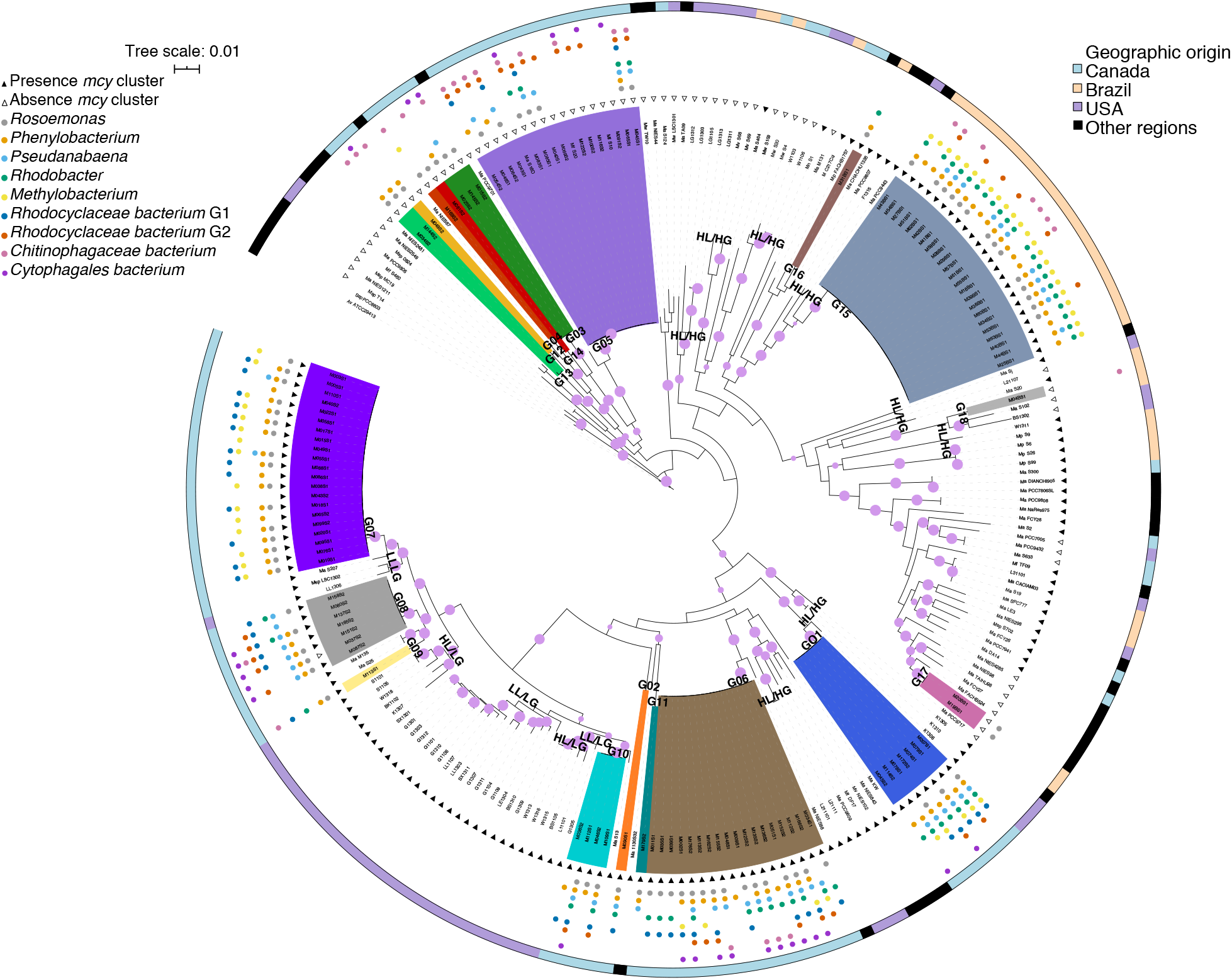
Maximum likelihood phylogenetic tree of 109 *Microcystis* colony genomes and previously sequenced reference genomes. *Microcystis* genomes were classified in 18 genotypes based on Average Nucleotide Identity (ANI) greater or equal to 99%. A core genome was inferred based on 109 *Microcystis* genomes and 122 *Microcystis* reference genomes downloaded from NCBI. The alignment of the 115 core genes (68,145 bp in total after excluding positions with gaps) was used to infer the Maximum Likelihood phylogeny. The tree was rooted using two cyanobacteria (*Anabaena variabilis* ATCC29413 and *Synechocystis* sp. PCC6803) as outgroups. The clades highlighted in different colours indicate *Microcystis* genotypes (G01 to G18) from this study; uncolored clades are other reference genomes from the literature. The purple circles on the tree branches indicate bootstrap values greater or equal to 70%. The empty and filled triangles around the tree indicate absence and presence of the *mcy* cluster, respectively. The small colored and filled dots indicate the most prevalent associated bacteria genera related to each *Microcystis* genome. The outermost circle indicates the geographic origin of the *Microcystis* genomes. Several references genomes of *Microcystis* genotypes recovered from environments with high- and low phosphorus are indicated as LL/LG (Low Phosphorus Lake/Low Phosphorus genotype), HL/LG (High Phosphorus Lake/Low Phosphorus Genotype) and HL/HG (High Phosphorus Lake/High Phosphorus Genotype).

*Microcystis* is thought to be adapted to high nutrient conditions, since it often blooms in eutrophic waters such as Champlain and Pampulha (Supplementary Table 3). However, a recent sampling of Michigan lakes identified *Microcystis* isolates adapted to low-phosphorus (low-phosphorus genotypes, LG), which occur in both high- and low-phosphorus lakes^30^. Genotypes G07, G08, G09 and G10 from Lake Champlain are nested within the LG clade with high bootstrap support (Fig. 1), indicating that low-phosphorus-adapted genotypes also occur in high-phosphorus lakes. Notably, most of the genomes within the LG clade (66 out of 67) encode the *mcy* gene cluster required for the biosynthesis of the cyanotoxin microcystin^32^. In contrast to the single LG clade, high-phosphorus genotypes (HG), are broadly distributed across the phylogenetic tree, recovered from multiple geographic locations, and some but not all encode *mcy* (Fig. 1). This pattern of *mcy* presence/absence is consistent with multiple *mcy* gene gain/loss events, mostly occurring in deep internal branches of the phylogeny, such that closely-related genotypes tend have identical *mcy* gene profiles.

### Lower *Microcystis* diversity within than between colonies of the same genotype suggests clonal colony formation

A previous study of Michigan lakes supported clonal colony formation (by cell division) in isolates from high-phosphorus lakes, but suggested a preponderance of nonclonal colonies (by agglomeration of distantly related cell) in low-phosphorus lakes^30^. To distinguish between clonal and nonclonal colony formation, we compared genetic diversity within and between colonies. Within colonies, the number of single nucleotide variants (SNVs) was significantly lower (mean of 3 SNVs) than between colonies (mean of 25) of the same genotype (Two-tailed Wilcoxon Rank Sum Test, *P* < 0.05; twelve outliers with more than 300 variants between colonies were excluded, making the test conservative) (Fig. 2 and Supplementary Table 4). These outliers were found in colonies within the genotypes G05, G06, G08 and G13. To put these results in context, *Microcystis* evolved an average of 5 SNVs after ~6 years of culture, slightly more variation than observed within a colony but still ~5X less than observed between colonies of the same genotype (Two-tailed Wilcoxon Rank Sum Test, *P* < 0.05). Overall, these results are consistent with colony formation occurring mainly by clonal cell division in Lake Champlain and Pampulha – at least under the sampled environmental conditions.

**Figure 2.**
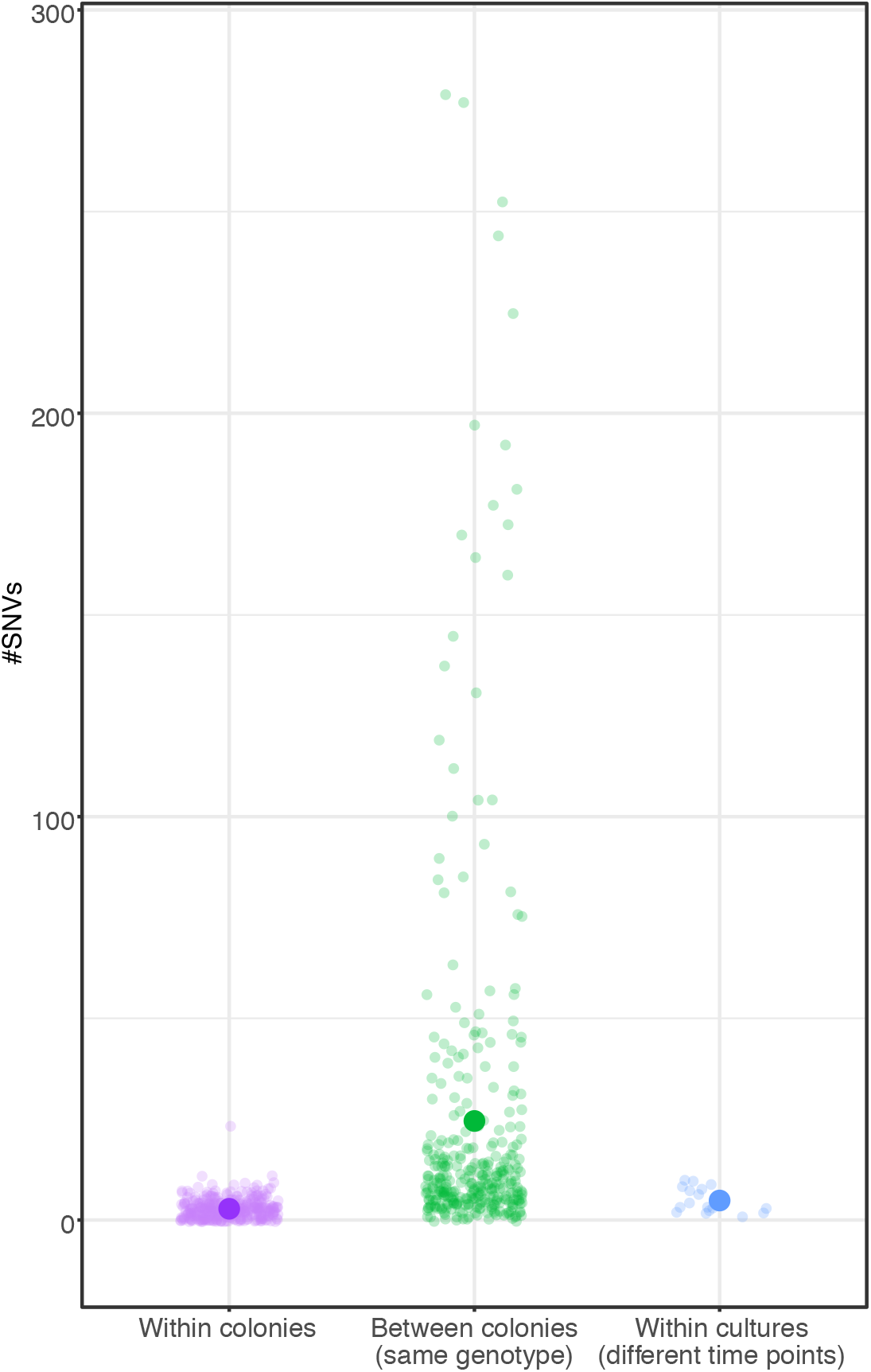
Greater genetic diversity between than within *Microcystis* colonies. The number of single nucleotide variants (SNVs) within and between *Microcystis* colonies of the same genotype are shown, compared to SNVs that occurred over ~6 years of *Microcystis* culture in the laboratory (Methods). Large points show mean values.

### Evidence for phylosymbiosis between *Microcystis* and its microbiome

Having characterized the genetic diversity of *Microcystis* genomes, we turned our attention to the colony-associated bacteria (AB). We recovered a total of 391 high-quality non-*Microcystis* genomes (Completeness ≥ 70 and contamination < 10%) from the 109 colonies (Supplementary Table 1 and 5), classified into 72 putative species (ANI > 95%) and 37 genera. Only five AB species were shared among colonies from Canada and Brazil: *Pseudanabaena* sp. A06, *Methylobacterium* sp. A30, *Roseomonas* sp. A21, *Burkholderia* sp. A55 (a likely contaminant, as discussed below) and *Gemmatimonas* sp. A63 (Supplementary Fig. 2). Because certain low-abundance AB might be present in a colony but fail to assemble into a high-quality genome, we mapped reads from each colony to a database of all the AB genome assemblies and estimated AB genome coverages; each colony contained an average of six AB (genome coverage greater or equal to 1X), with a range of 0 to 15 (Supplementary Fig. 3). We found no strict “core” of AB present in all colonies, either at the species or genus level. However, several genera were quite prevalent. These include *Phenylobacterium* (present in 73.40% of colonies), *Roseomonas* (70.64%), *Pseudanabaena* (43.12%), *Rhodobacter* (46.79%), *Methylobacterium* (44.04%), *Rhodocyclaceae* G1 (unclassified genus) (39.45%), *Rhodocyclaceae* G2 (unclassified genus) (31.19%), *Chitinophagaceae* (unclassified genus) (26.60%) and *Cytophagales* (unclassified genus) (22.94%).

To assess the evidence for phylosymbiosis, we first asked if different *Microcystis* genotypes had distinct colony microbiomes. The phylogeny illustrates how certain *Microcystis* genotypes appeared to be preferentially associated with particular AB (Fig. 1). For example, *Phenylobacterium* and *Methylobacterium* were present in all the colonies of genotype G15, while *Rhodobacter* and *Phenylobacterium* occur in all colonies of genotype G01. These anecdotal patterns are borne out in analyses of colony community structure, which show that *Microcystis* genotypes have distinct microbiomes (Fig. 3a). Genotype explains more variation in community structure (PERMANOVA on Bray-Curtis distances, *R*^*2*^ = 0.387, *P* < 0.01; Supplementary Table 6) than any other measured variable including pH (*R*^*2*^ < 0.05) or temperature at the sampling site (*R*^*2*^ < 0.05), presence of microcystin (*mcy*) genes in the genotype (*R*^*2*^< 0.05), or sampling site (*R*^*2*^ = 0.11). Genotype was still the best explanatory variable when the analysis was performed on Lake Champlain samples only (Fig. 3b, PERMANOVA, *R*^*2*^ = 0.309, *P* = 0.001). A key piece of evidence for phylosymbiosis is not only for microbiomes to differ among host lineages, but for microbiome composition to change proportionally to host phylogeny. To test this, we converted the *Microcystis* host phylogeny into a distance matrix, which we correlated with the colony microbiome Bray-Curtis dissimilarity matrix. Consistent with phylosymbiosis, we found that microbiome composition changes were correlated with the host phylogeny according to a Mantel test (*r* = 0.5, *P* = 0.001) confirmed with Procrustean superimposition (*r* = 0.6, *P* = 0.001)^33^.

**Figure 3.**
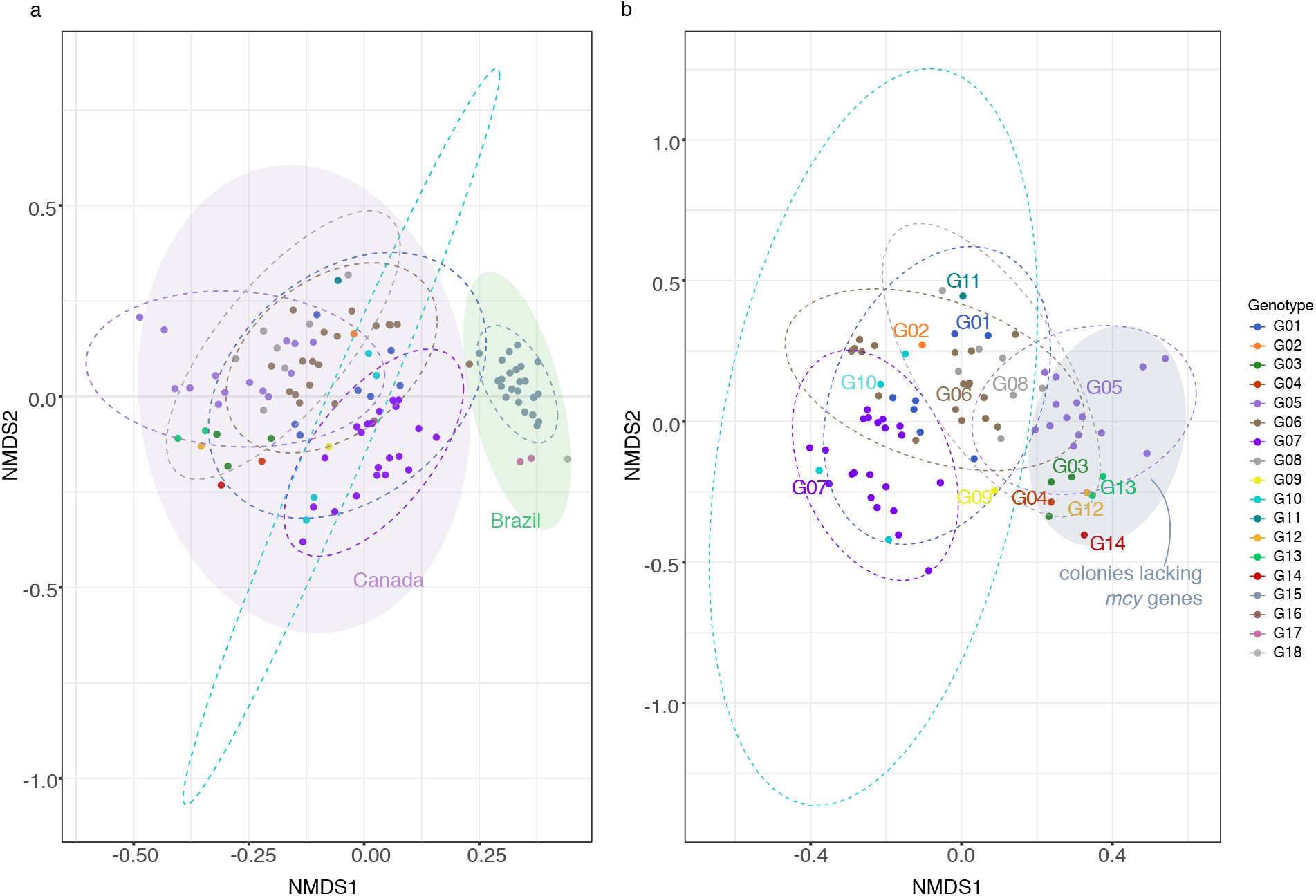
*Microcystis* genotypes have distinct microbiomes. Non-metric multidimensional scaling (NMDS) plots are based on the coverage of the non-*Microcystis* metagenome-assembled genomes (MAGs) per colony (Bray–Curtis distance). **a)** All samples, including those from Pampulha, Brazil and Lake Champlain, Canada. Ellipses show 95% confidence intervals (stress = 0.202). **b)** Samples from Lake Champlain only (stress = 0.225). The grey shaded ellipse shows *Microcystis* colonies that do not encode the *mcy* cluster for microcystin toxin production.

### *Microcystis* genotype abundances vary over time in Lake Champlain and are correlated with prevalent members of the microbiome

*Microcystis* producers and non-producers of the cyanotoxin microcystin are known to change in relative abundance within lakes over time^31,34,35^. More generally, to what extent different genotypes of *Microcystis* vary over time, along with their colony-associated bacteria, is less well known. We investigated the *Microcystis* genotype diversity in metagenomes from Lake Champlain based on 14 *Microcystis* genotypes identified in colonies from 2017 and 2018 (Fig. 1). Using a gene marker database of these 14 *Microcystis* genotypes (Methods), we estimated the relative abundance and read coverage of each genotype in 72 metagenomes from 2006 to 2018, sampled during the summer months (Supplementary Fig. 4). It is possible that these 14 genotypes do not represent the total genotypic diversity of *Microcystis* occurring in the lake. However, mapping metagenomic reads from the lake to these genotypes with a 99% sequence identity threshold allowed us to recover 93.5% of *Microcystis* reads (defining *Microcystis* at 96% sequence identity).

Using a distance-based redundancy analysis (dbRDA), we estimated the effect of total phosphorous, total nitrogen, dissolve phosphorous, dissolved nitrogen, mean temperature and time (years, months and season) on the *Microcystis* genotype community composition in the 42 Lake Champlain metagenomes with complete metadata, and with *Microcystis* genome coverage greater or equal to 1X. *Microcystis* genotype diversity in environmental metagenomes was best explained by yearly temporal variation (*R*^*2*^ = 0.511, *P* = 0.002; Supplementary Fig. 5). Years did not differ significantly in their dispersion (PERMDISP *P* > 0.05; Supplementary Table 6). Environmental variables such as nitrogen and phosphorus did not have a significant effect on the community composition. In a shorter time series (April to November of one year) in Pampulha, a more diverse community of four *Microcystis* genotypes eventually came to be dominated by one genotype (G15) encoding the *mcy* toxin biosynthesis gene cluster (Supplementary Fig. 6). However, more extensive sampling is required to estimate the effect of other environmental variables (i.e., phosphorus) on the community composition in Brazil.

Similarly to *Microcystis* genotypes, the composition of AB in Lake Champlain also varied significantly across years (PERMANOVA, on Bray-Curtis distances, *R*^*2*^ = 0.43, *P* < 0.01; Supplementary Fig. 7, stress = 0.1569). We asked if the presence of dominant *Microcystis* genotypes could explain the variation in the AB community composition. A significant effect of the genotype was observed using PERMANOVA (*R*^*2*^ = 0.14, *P* < 0.01), but not using dbRDA (*R*^*2*^ = 1.2, *P* > 0.05). Years and *Microcystis* genotypes were the best explanatory variables for AB composition; however, their dispersions were significantly different (*P* < 0.01) making the PERMANOVA results difficult to interpret. In addition, the AB community sampled from metagenomes includes both free-living and colony-attached AB, possibly adding noise to any signal of *Microcystis* genotypes selecting for specific AB within colonies.

We further hypothesized that the most prevalent AB in *Microcystis* microbiome should co-occur with *Microcystis* in lake metagenomes. In contrast, they should not co-occur with another cyanobacterium frequently observed in Lake Champlain, *Dolichospermum*, which serves as a negative control. We first estimated normalized read counts and coverage of *Microcystis*, *Dolichospermum* in the 72 metagenomes from the Lake Champlain time series (Supplementary Fig. 8). We then estimated the Spearman correlations between *Microcystis* or *Dolichospermum* and each AB species or genus. The two cyanobacteria were weakly correlated across the environmental metagenomes (*r* = 0.29 and *Q-value* = 0.027, Spearman rank-based correlation test). As expected, the nine most prevalent AB genera in the *Microcystis* microbiome were strongly correlated with *Microcystis* (*r* > 0.7, *Q-value* < 0.001), and only weakly with *Dolichospermum* (*r* < 0.4, *Q-value* > 0.001) with the exception of *Phenylobacterium* (*r* = 0.47, *Q-value* < 0.001) which is nevertheless more strongly associated with *Microcystis* (Supplementary Fig. 9). The positive correlation between the most prevalent AB genera and *Microcystis* was also supported using an alternative correlation method, SparCC, which corrects for compositional effects in the data (*r* > 0.4, *Q-value* < 0.05) (Supplementary Table 7 and Fig. 9c). These significant positive correlations are consistent with close interaction between *Microcystis* and the most prevalent genera related to their microbiome. Genera found at lower prevalence in *Microcystis* colonies (*e.g., Phycisphaerales bacterium* (unclassified genus) and *Telmatospirillum*) were poorly correlated with both *Microcystis* and *Dolichospermum* (Supplementary Table 7 and Fig. 9a). Another AB belonging to the genus *Burkholderia* was quite prevalent in colonies but poorly correlated with *Microcystis* in metagenomes (present in the 40.37% of the colonies; *r* = −0.16, *Q-value* = 0.343) suggesting likely contamination of colonies rather than a true ecological association. However, such a signal of contamination was rare, suggesting that most of the data reflect true associations.

Finally, we asked if specific *Microcystis* genotypes were correlated with the presence of specific AB species (Supplementary Fig. 10) observed in *Microcystis* colonies. For example, *Rhodocyclaceae bacterium* G2 A13 was better correlated with genotype G05 than other *Microcystis* genotypes, consistent with the prevalence of this species in 13 out of 14 colonies of genotype G05. In contrast, genotype G10 was poorly correlated with certain species within the genera *Roseomonas* and *Methylobacterium* (r < 0.38, *Q-value* > 0.001). Overall, this is consistent with certain *Microcystis* genotypes having strong preferences for certain AB, while being unselective for others.

### Signatures of co-speciation between *Microcystis* and members of its microbiome

Phylosymbiosis can arise due to vertical inheritance of microbiomes, or horizontal acquisition of microbiomes at each generation, provided that host lineages are matched with distinct microbiomes. To assess the evidence for vertical inheritance of *Microcystis* AB, we used ParaFit to test for similarity between the *Microcystis* phylogeny and the phylogenies of the nine most prevalent AB genera strongly correlated with *Microcystis* but not with *Dolichospermum* in Lake Champlain (Supplementary Fig. 9). Each of these genera was represented by at least 12 high-quality draft genomes and was found in at least five different *Microcystis* genotypes. Significant co-phylogenetic signal suggests co-speciation of hosts and symbionts, consistent with a relatively long evolutionary history of association (*e.g.*, vertical descent). We found that *Roseomonas*, the second most prevalent AB genus in colonies, and *Rhodobacter*, the third most prevalent, had significant signatures of co-phylogeny (Fig. 4), while *Phenylobacterium* and *Chitinophagaceae* were borderline cases (Table 1). Overall, there was no clear tendency for stronger co-phylogeny with more prevalent AB, or with AB most correlated with *Microcystis* over time in Lake Champlain metagenomes (Table 1). However, such tendencies would be hard to discern in this relatively small sample size. As expected, the likely contaminant *Burkholderia* A55 (*Burkholderia cepacia*) present in 40.37% of colonies, was poorly correlated with the presence of *Microcystis* in environmental metagenomes (*r* = −0.16, *Q-value* = 0.343), with no signal of co-phylogeny (*P-value* = 0.732). Although co-phylogenetic signal was detectable in at least two of the most prevalent AB, the phylogenies are not identical (Fig. 4), suggesting a mixture of vertical and horizontal transmission. Even if horizontal transmission of AB among *Microcystis* lineages is likely, some degree of host-microbiome matching must be occurring to explain the co-phylogenetic signal.

**Figure 4.**
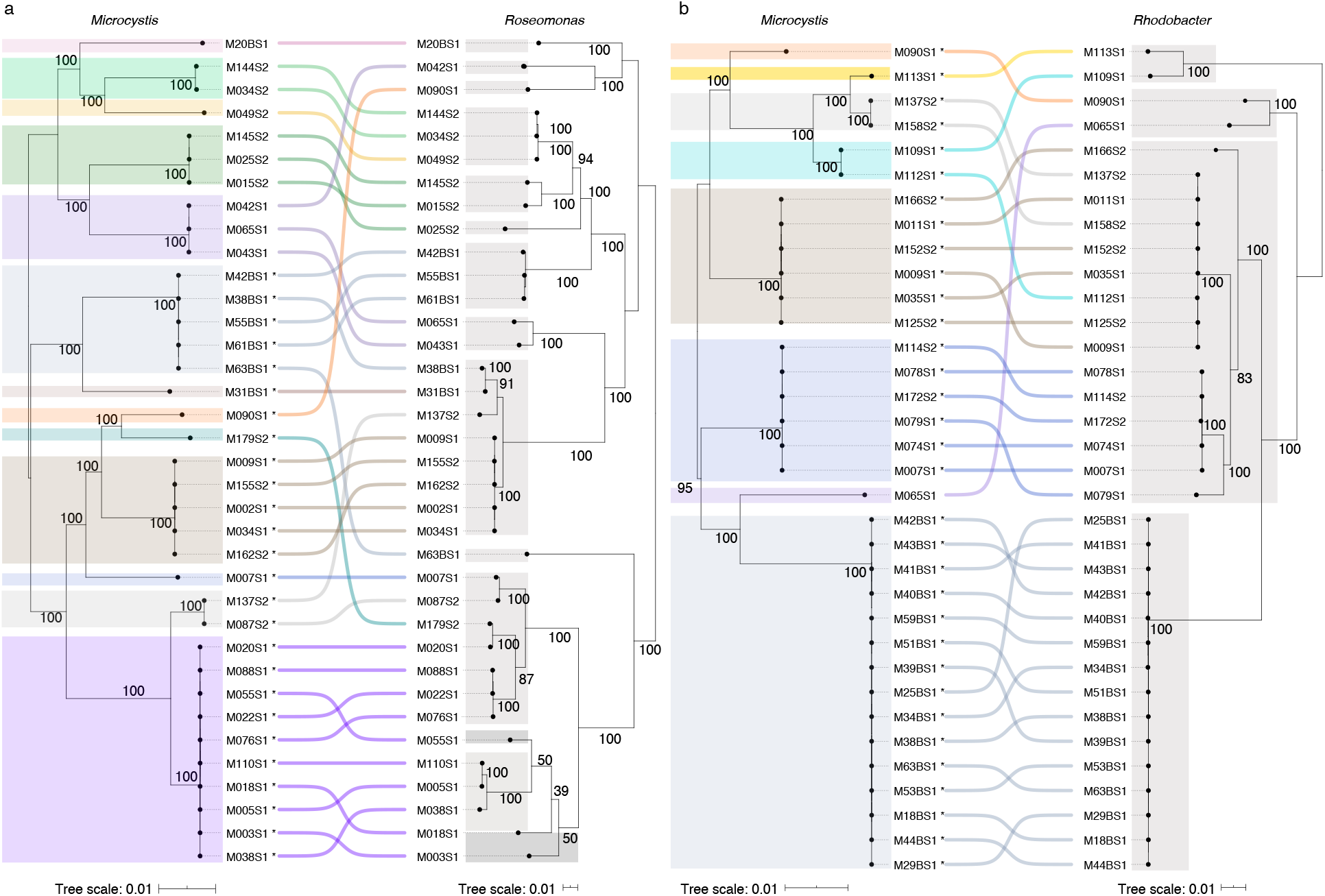
Co-phylogeny between *Microcystis* and two prevalent associated bacteria. (a) *Roseomonas* and (b) *Rhodobacter* core genome phylogenies were compared to the *Microcystis* core phylogeny. The lines between the two phylogenies connect genomes coming from the same *Microcystis* colony. The phylogenetic trees for *Microcystis*, *Roseomonas* and *Rhodobacter* were based on 706, 135 and 470 core genes, respectively. The different *Microcystis* genotypes are highlighted in colour, and the *Roseomonas* or *Rhodobacter* species in gray. The asterisks indicate the presence of the *mcy* cluster. The co-phylogenetic similarity is greater than expected by chance (ParaFit Global test, *P-value* < 0.01).

### Horizontal gene transfer (HGT) between *Microcystis* and its associated bacteria

Unrelated bacteria sharing a common environment, such as the human gut, are known to engage in frequent horizontal gene transfer^36^. We hypothesized that *Microcystis* would also exchange genes with members of its microbiome, which share a similar ecological niche – the colony milieu – for at least some period of time. We began by using a simple heuristic to look for similar gene sequences (≥ 99% amino acid identity) occurring in the *Microcystis* genome and at least one AB genome, as a proxy for relatively recent HGT events. Genome assembly and binning could affect this analysis by misplacing identical sequences either in *Microcystis* or in an AB genome, but not in both. To reduce this bias, we only considered a gene to be involved in HGT if it was present in at least four genomes. We identified a total of 1909 genes involved in HGT between *Microcystis* and one of seven AB species: *Pseudanabaena* A06, *Pseudanabaena* A07*, Burkholderiales bacterium* G3 A12, *Rhodocyclaceae bacterium* G2 A13, *Chitinophagaceae bacterium* A08, *Cytophagales bacterium* A04 and *Cytophagales bacterium* A05. Compared to the *Microcystis* core genome, these genes are enriched in functions related to secondary metabolite biosynthesis, replication and recombination, and defense mechanisms (Fig. 5). As a control, we repeated the analysis of HGT using the likely contaminant *Burkholderia* A55 genome instead of *Microcystis*. We identified 558 putative HGT events, of which 523 involving species not found to engage in HGT with *Microcystis: Methylobacterium* A30, *Rhodocyclaceae bacterium* G1 A54 and *Cupriavidus* A44. This suggests that *Microcystis* engages in more HGT with its microbiome than a random expectation (*i.e.* with a contaminant genome), and allows us to conservatively estimate the false-positive rate of HGT detection at 523/(523+1909), or 22%. Despite the significant noise, we expect the broad gene functional categories and specific AB involved in HGT with *Microcystis* to be relatively robust (Fig. 5). Surprisingly, prevalent AB with evidence of co-phylogeny with *Microcystis* (*Roseomonas* and *Rhodobacter*) shared relatively few (less than seven) HGT events with *Microcystis*. This counter-intuitive result could be explained if these co-phylogenetic associations are relatively ancient, but our HGT detection is biased toward recent events. Alternatively, it is possible that HGT is more likely among less intimately associated bacteria, whereas an intimate association would select for only one, but not both partners, to encode the gene. This would also require that metabolites are shared between partners. Further work will be needed to thoroughly test this hypothesis.

**Figure 5.**
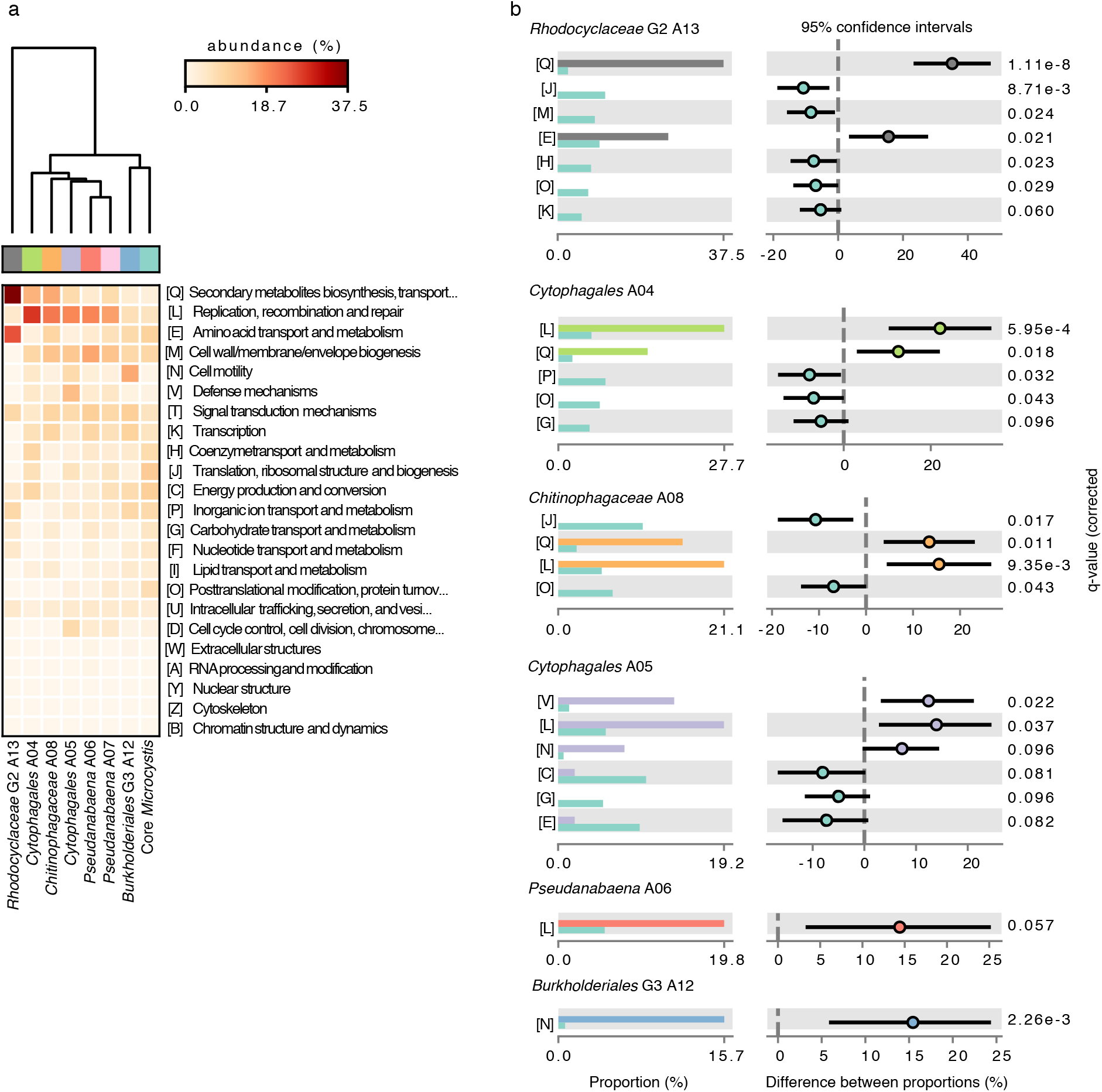
Inferred recent HGT between *Microcystis* and associated bacteria. Horizontal transferred genes between *Microcystis* and each AB species were inferred with a simple heuristic and annotated in 23 Clusters of Orthologous Groups (COGs) functional categories using EggNOG mapper (Methods). **a)** Clustering analysis based on the relative abundance of the genes for each functional category, compared to the genes in the *Microcystis* core genome. **b)** COG functions showing differential abundance between *Microcystis* core genes (turquoise) and the set of putative HGTs (other colors).

As an additional validation of our HGT heuristic, we used Metachip, which uses phylogenetic incongruence in addition to a sequence identity threshold^37^. Metachip identified the same seven AB genera involved in HGT with *Microcysis* based on our simple heuristic, except for *Rhodocyclaceae bacterium* G2. However, Metachip is much more conservative, identifying only 46 gene families involved in HGT (Supplementary Table 8). Of these gene families 31 were also identified by our heuristic method, suggesting they are high-quality candidates.

### Cellular functions encoded by members of the *Microcystis* microbiome

In contrast to genes shared by HGT, there may be a genetic division of labour between *Microcystis* and its microbiome, which would then be expected to encode different and complementary sets of gene functions. To compare these gene functions, we first characterized orthologous genes using the Kyoto Encyclopedia of Genes and Genomes (KEGG) orthologues (KO) in both *Microcystis* and its microbiome. Then, using the software ANTISMASH, we identified gene clusters involved in the biosynthesis of cyanopeptides and other pathways of interest. As expected for distantly related bacteria, *Microcystis* genotypes and AB encode distinct sets of gene functions (Supplementary Fig. 11). Bacteria from the same Phylum tend to cluster together in terms of their functional gene content. For example, *Microcystis* genotypes clusters with its fellow cyanobacteria *Pseudanabaena*, while Bacteroidetes (*i.e. Cytophagales bacterium* and *Chitinophagaceae bacterium*) formed a distinct cluster (Supplementary Fig. 11).

We identified several examples of possible functional complementarity between *Microcystis* and members of its microbiome. For example, *Microcystis* encodes incomplete pathways for the synthesis of biotin (M00123; pimeloyl-ACP/CoA => biotin) and cobalamin (M00122; cobinamide => cobalamin), suggesting that these functions might be subject to gene loss if the functions are provided by the microbiome. Consistent with this idea, AB encode complete pathways for both biotin (in *Cytophagales*, *Chitinophagaceae* and *Rhodocyclaceae*) and cobalamin (in *Rhodobacter, Azospirillum*, and *Bradyrhizobium*). Other AB (*e.g.*, *Roseomona*s, *Rhodobacter* and *Methylobacterium*) encoded genes involved in the anoxygenic photosynthesis (Supplementary Table 9) and genes related with the transport of rhamnose, D-xylose, fructose, glycerol and a-glucoside, which could also complement the metabolic repertoire of *Microcystis*^16^, although this deserves further study.

*Roseomonas* and *Rhodobacter*, which show co-phylogeny with *Microcystis* but appear not to engage in significant amounts of HGT, are prime candidates for functional complementarity to have evolved and be maintained with high partner fidelity. Both these genera encode genes for the biosynthesis of carotenoids (phytoene desaturase (*crtI*) and phytoene synthase (*crtB*)). Carotenoid pigments like zeaxanthin are generally produced by *Microcystis* for their photoprotective properties and their capacity to improve the efficiency of photosynthesis.^38^ Indeed, in our *Microcystis* genomes, we found genes encoding for phytoene synthase (*crtB*) and zeaxanthin glucosyltransferase (*crtX*). However, genes like (*crtI*), lycopene cyclase (*crtY*) and beta-carotene hydroxylase (*crtZ*) were only found in other AB genomes (*e.g*., *Cytophagales*). It is tempting to speculate that the *Microcystis* microbiome may also be involved in the production of these carotenoids. *Roseomonas* and *Rhodobacte*r also have metabolic pathways for nitrogen fixation. *Microcystis* is unable to fix nitrogen, and previous studies have suggested it may rely on its microbiome for nitrogen^16,39^. The co-phylogenetic signal between *Microcystis* and these genera might thus be explained by these complementary functions.

## Discussion

By combining single colony sequencing and metagenome analysis, we explored the genetic diversity of both *Microcystis* and its microbiome, and their variation over time in Lake Champlain, Canada and the Pampulha reservoir in Brazil. We revealed a higher diversity of *Microcystis* genotypes than previously described^40^, and patterns of cophylogeny, phylosymbiosis and HGT between the host and its microbiome. Despite the absence of a core microbiome, several of the associations between *Microcystis* and its attached bacteria, notably *Roseomonas* and *Rhodobacte*r, appear to be relatively stable over evolutionary time. These two genera have been previously reported to be correlated with *Microcystis* in environmental samples^41,42^. Whether these associations are beneficial to one or both partners remain to be seen, and deserve further study as possible targets for better predicting and controlling harmful *Microcystis* bloom events. For example, small filamentous cyanobacteria *Pseudanabaena* and members of the order *Cytophagales* have been previously reported as bloom biomarkers^43^.

There has been some debate about whether *Microcystis* colonies form by clonal cell division, or by aggregation of (potentially distantly related) cyanobacterial cells^21,44^. Consistent with another recent study in eutrophic lakes^30^, we conclude that clonal cell division is more likely, based on our observation of much greater genetic variation in the *Microcystis* genome between than within colonies of the same genotype. One caveat to this conclusion is that our limited and possibly biased sample of *Microcystis* colonies means that aggregated colonies could exist, but were unsampled due to small colony size (resulting in failure of DNA extraction). However, 93.5% of *Microcystis* metagenomic reads from Lake Champlain were recruited to our collection of colony genomes at 99% nucleotide sequence identity, suggesting that the majority of natural *Microcystis* diversity is represented in our sample of colonies. Of course, these results are specific to Lake Champlain and should be replicated in other lakes under different environmental conditions (*e.g.*, oligotrophic lakes).

Phylosymbiosis and co-speciation appear to be relatively common and strong in mammalian gut microbiomes^22,23^, and even in the more environmentally-exposed coral microbiome^22,23^. It is unclear if such tight and evolutionarily stable associations would apply to *Microcystis* and its associated bacteria, or if more transient interactions would prevail. While the idea of a *Microcystis* microbiome has been suggested previously based on bulk metagenomic and amplicon sequencing from lakes ^16,45^, here we refine the *Microcystis* microbiome concept beyond co-occurrence patterns to physical association within a colony. We found that the most prevalent associated bacteria from individual *Microcystis* colonies also tend to co-occur with *Microcystis* over time in Lake Champlain. The composition of the microbiome varies along the *Microcystis* phylogenetic tree, consistent with phylosymbiosis and relatively long-term associations. At least two associated bacteria show significant co-phylogenetic signal, suggesting co-speciation with *Microcystis.* Therefore, although possibly not as strong as in mammals or even coral, phylosymbiosis and co-phylogeny are features of the *Microcystis* microbiome. Phylosymbiosis can arise as a consequence of shared biogeography between hosts and microbiomes^46^, and we do observe distinct microbiomes in Brazil and Canada. However, we found evidence for phylosymbiosis within a single lake in Canada, suggesting that other factors – such as host-microbiome trait matching – are likely at play.

As expected for distantly related bacteria, *Microcystis* and its associated bacteria encode different functional gene repertoires, some of which could be complementary and mutually beneficial. For example, we found that associated bacteria may complement biosynthetic functions that were lost or never present in *Microcystis*, such as biotin, cobalamin, or carotenoid synthesis. Carotenoids act as antioxidants and may increase the photosynthetic light absorption spectrum^47,48^. Some associated bacteria, including the co-speciating *Roseomonas* and *Rhodobacte*r, have metabolic pathways for nitrogen fixation and phosphonate transport. *Microcystis* is unable to fix nitrogen, and studies suggest that it may rely on nitrogen-fixing members of its microbiota^16,39^. While it remains unclear if metabolites are actually exchanged between *Microcystis* and members of its microbiome, these hypotheses could be tested experimentally.

Horizontal gene transfer (HGT) is relatively common in bacteria, and may occur among unrelated bacteria^49^ particularly when they share an ecological niche such as the human gut^36^. *Microcystis* is physically associated with its microbiome for at least part of the colony life cycle, and we hypothesized that HGT could occur within colonies. Using two methods to detect HGT, we found evidence for gene transfers between *Microcystis* and at least six different species of associated bacteria: two species of *Pseudanabaena*, two *Cytophagales*, one *Burkholderiales*, and one *Chitinophagaceae* species. Notably, we did not find evidence for HGT between *Microcystis* and its two most co-phylogenetically associated bacteria, *Roseomonas* and *Rhodobacte*r. To explain this result, we hypothesize that such long-term associations might favour the loss of redundant genes, as predicted by the Black Queen Hypothesis^50^. In other words, a gene needs to be encoded by only one partner, provided that gene products or metabolites are shared between partners. Therefore, even if HGT does occur between partners, we would not expect to find the same gene redundantly encoded in both partners. These evolved co-dependencies would further reinforce partner fidelity and could help explain the co-phylogenetic signal between them.

Overall, our results provide evidence for long-lasting eco-evolutionary associations between *Microcystis* and its microbiome. Some members of the microbiome may be more tightly associated than others, and based on their gene content we hypothesize that they may provide beneficial and complementary functions to *Microcystis.* These hypotheses could be tested in experimental co-cultures, which have recently shown how the *Microcystis* microbiome can alter its competitive fitness against eukaryotic algae^51^. These experiments could be extended to the combinations of *Microcystis* genotypes and associated bacteria which we have shown to be intimately associated in nature.

## Methods

### Sample collection and DNA extraction for colonies and metagenomes

To access to the genomic diversity of *Microcystis* in Lake Champlain and Pampulha reservoir, 346 individual *Microcystis* colonies were isolated across the bloom season (July to October in Quebec, Canada (45°02′44.86′′N, 73°07′57.60′′W) and April to November in Minas Gerais, Brazil (19°55′09″S and 43°56′47″W)). Colonies were isolated from surface water samples (~50 cm depth) after concentration using a plankton net (mesh size 20 μm). One liter of concentrated water was collected and stored at 4 °C for a maximum of 36 hours until colony isolation. Colonies were isolated using micropipes, sterile medium (Z8 medium) and a microscope (Nikon E200 Eclipse). Each colony was washed 15-20 times using sterile Z8 medium and stored at −80 C until DNA extraction. The DNA extraction was performed directly on each colony using the ChargeSwitch® gDNA Mini Bacteria Kit. Two additional steps were added to ensure the rupture of the *Microcystis* colonies and cells (See Supplementary Methods). Briefly, each colony was added to a tube containing 50 mg of beads (PowerBead tubes, glass 0.1 mm-Mo-bio), incubated with lysis solutions, and then vortexed using the TissueLyser LT (Qiagen) for three minutes at 45 oscillation per second. The tube was then centrifuged for 1 minute at 9000 rcf. This procedure yielded DNA for 109 colonies, sequenced as described below. Matched water samples were collected at the same place and time as colonies, spanning 16 time points (Supplementary Table 10). Water temperature and pH were also measured at each sampling point.

For metagenomic sequencing, a total of 72 lake water samples were collected over 10 years (2006 to 2018) during the ice-free season (April to November) from the photic zone of Missisquoi Bay at two different sites (littoral and pelagic) of Lake Champlain, Quebec, Canada (45°02′45′′N, 73°07′58′′W). Lake water was filtered and DNA was extracted using a Zymo Kit (Zymo, D4023) as described previously^43^. The filtration was performed the same day of the sampling, using between 50 and 250 mL of water samples, depending on the amount of biomass, onto 0.2 μm hydrophilic polyethersulfone membranes (Millipore, Etobicoke, ON). Samples were obtained at relatively low frequency between 2006 and 2016, and at higher frequency (approximately weekly or more often) during bloom periods between 2015 and 2016 (Supplementary Table 3). Water samples corresponding to six sampling points from Minas Gerais Brazil were also collected for DNA extraction and metagenome sequencing. Environmental variables were measured for each sample. Sample water were collected (50 ml) for measuring nutrients (DN, DP, TP and TN), except for the samples from Brazil (Supplementary Table 3)^43^.

### DNA sequencing of single colonies and metagenomes

DNA extracted from *Microcystis* single colonies was sequenced using the Illumina HiSeq 4000 platform with 150bp paired-end reads. The sequencing libraries (with average fragment size 360bp) were prepared using the NEB (New England Biolabs®) low input protocol. The DNA extracted from filtered bulk lake water for each sampling point (2017 and 2018) from Canada and Brazil were sequenced using Illumina NovaSeq 6000 S4 platform with 150bp paired-end reads. The earlier lake water samples from a previous long-term experiment in Lake Champlain (2006 to 2016) were sequenced using Illumina Hiseq2500 with 125 paired-end reads (Supplementary Table 3).

### Metagenome assembly and genome binning

For the *Microcystis* colonies, the sequencing reads were filtered and trimmed using Trimmomatic (v0.36)^52^ then assembled with MEGA-HIT (v1.1.1)^53^, producing contigs belonging to both *Microcystis* and associated bacteria. We then used Anvi’o (v3.5) to filter, cluster and bin the contigs longer than 2,500 bp as was previously^29,54^. The quality of each resulting metagenome-assembled genome (MAG) was estimated using CheckM (v1.0.13)^55^. From the 109 colonies, 500 medium and high-quality MAGS were identified (completeness ≥ 70% and contamination ≤ 10%) (Supplementary Table 1 and 5)^56^. MAGs were annotated using Prokka (v1.14.0)^57^. Pairwise average nucleotide identity (ANI) values between genomes were estimated using FastANI (v1.2) and pyani^58,59^. MAGs were classified into different taxonomic groups at a threshold of ANI ≥ 96% (Supplementary Table 5 and 11). MAGs were assigned to genera and species using Blastp of the recA and RpoB proteins against the NCBI database, and refined using the Genome Taxonomy Database Toolkit (GTDB-Tk) (v1.0.2), which uses a set 120 universal bacterial gene markers^60^.

For each taxonomic group, we selected at least two representative sequence types (for a total of 138 genomes), from which we inferred a Maximum likelihood phylogenetic tree based on the core gene alignment using RAxML (v8.2.11)^61^. The core genome was estimated using panX (v1.5.1). Core genes were defined as those genes present in at least the 80% of sampled genomes (e-value < 0.005)^62^. Each of the resulting 62 core genes was alignment using muscle (v3.8.3)^63^. Filter.seqs from mothur (v1.41.3) was used to remove the gaps per each gene alignment^64^. Individual alignments were concatenated into a single alignment (16,400 bp long) input into RAxML.

### Assessment of the *Microcystis* genotype diversity in freshwater colonies

A core genome was also estimated for the 109 *Microcystis* genomes and 122 NCBI references genomes (Supplementary Table 1 and 12). The resulting alignment of the 115 core genes was degaped (68,145 bp long) and used to infer an ML phylogeny using RAxML. Two outgroups (*Anabaena variabilis* ATCC29413 and *Synechocystis* sp. PCC6803) were included. Based on ANI values greater or equal to 99%, the monophyletic clades of *Microcystis* genomes were classified into 18 genotypes (Supplementary Table 2).

### Assessment of the *Microcystis* genomic (within-colonies) variation versus intra-genotype variation (between colonies)

We first confirmed that *Microcystis* is haploid, as polyploidy has been observed among other cyanobacteria^65^. We estimated ploidy variation in *Microcystis* colonies using k-mer frequencies and raw sequences. We first mapped the reads of each colony (containing reads from both *Microcystis* and its microbiome) to a *Microcystis* reference genome using BBmap with minimum nucleotide identity of 99%^66^. Mapped reads were extracted using Picard (http://broadinstitute.github.io/picard/) and analyzed using Genomescope and Smudgeplot (https://github.com/tbenavi1/genomescope2.0; https://github.com/KamilSJaron/smudgeplot). All colonies appeared to be haploid, with a low rate of heterozygosity that could be due paralogs.

To determine whether *Microcystis* colonies likely formed by clonal cell division or cell aggregation, we called single nucleotide variants (SNVs) within colonies and between colonies of the same genotype. As a point of comparison, we also called SNVs that occurred over a period of approximately six years in laboratory cultures of *Microcystis* with genome sequences reported previously^29^. We used snippy (v4.4.0) (https://github.com/tseemann/snippy) with default parameters to call SNVs. Genotypes represented by only one sampled colony were excluded from the analysis (G02, G04, G09, G11, G12, G16, and G18).

SNV calling within and between colonies was executed by mapping reads against reference genomes. This was done independently for each genotype. We selected at least four reference genomes per genotype when possible. SNVs within colonies were detected by mapping the reads of the references to their respective genome assemblies. SNVs between colonies were detected by mapping the reads of different colonies of the same genotype to the genome assemblies of the references. We ignored positions where the reference nucleotide was poorly supported (threshold percentage for the minor variant <14.4%; mean = 1.1%) by the reads in both the within- and between-colony read mapping analyses because these were considered to be assembly errors.

### Identifying associated bacterial genomes in colonies

Non-*Microcystis* MAGs from each colony were classified in 72 species based on taxonomical analysis and ANI values ≥ 96%. Because individual assemblies could affect MAG completeness, we created a custom database of the 59 associated bacterial genomes from Quebec, and another database for the 18 species from Brazil. Using MIDAS (v1.3.0)^67^, we mapped the reads from each colony (downsampled to 8,000,000 reads per colony) against the custom databases to estimate the relative abundance and coverage for each of the 72 associated bacterial species. We defined a species to be present when it had a genome-wide average coverage of 1X or more. This allowed us to generate a matrix of associated bacteria presence or absence across colonies.

### *Microcystis’* microbiome composition variation according to environmental variables and host genotype

We first performed a distance-based RDA with the square root of the Bray-Curtis distance from a coverage table describing the composition of the *Microcystis* microbiome for each genotype. The variables included genotype information, presence/absence of *mcy* genes, temperature, pH, site (Canada or Brazil) and the temporal variables years and months. In a second approach, we calculated the beta diversity using the same dissimilarity distance and tested *Microcystis* microbiome composition variation using adonis() and betadisper().

We quantified phylosymbiosis by comparing the phylogenetic distance matrix of *Microcystis* genotypes and the microbiome composition distance matrix using a Mantel test (999 permutations, Spearman correlation) and the protest() R function to test the non-randomness between these two matrices (999 permutations) (vegan R package). The pairwise phylogenetic distances matrix was estimated using the RAxML tree of the *Microcystis* core genome and the cophenetic.phylo function of the ape R-package (v5.3)^68^.

### *Microcystis* genotypic diversity from metagenomic samples

*Microcystis* genomes from Quebec and Brazil were classified into 14 and four genotypes, respectively. This genotype classification was based on pairwise genome similarities greater or equal to 99%. Using the *Microcystis* genotypes and the software MIDAS (v1.3.0)^67^, we built two custom gene marker databases for the *Microcystis* genotypes (15 universal single-copy gene families), one for genotypes from Quebec and the other for genotypes from Brazil.

Using MIDAS and the custom databases, we estimated the relative abundances, the read counts and the read coverage of the *Microcystis* genotypes in 72 shotgun metagenomes from Lake Champlain, Quebec (62 metagenomes from a long-term experiment (2006 to 2016, excluding 2007 and 2014), plus 10 metagenomes from 2017 and 2018). Due the low number of *Microcystis* genotypes and metagenomes (6 sampling points for Brazil during 2018) from Brazil, these samples were not formally analyzed. Metagenomic reads with similarity greater or equal to 99% were mapped against the MIDAS database of *Microcystis* genotypes. We used 14,000,000 reads per metagenome after downsampling to the lowest-coverage metagenome (Supplementary Table 3). The metagenome sequencing from Brazil were mapped against a separate MIDAS database of the four *Microcystis* genotypes from Brazil (Supplementary Fig. 12).

To test if the 14 *Microcystis* genotypes represented in the colony genomes representative of the diversity present in the Lake Champlain metagenomes, we first mapped the downsampled metagenomic reads to a custom database including a single reference *Microcystis* genome (M083S1) (alignment identity cutoff = 96%), and also mapped the reads to the database including all the 14 genotypes (alignment identity cutoff = 99%). By using a cutoff value equal to 96%, we expect to recover most sequences from the *Microcystis* genus, regardless of which genotype the reads come from. We recovered 102,608 reads at 99% identity and 109,729 at 96%, showing that the 14 genotypes (defined at 99% identity) account for 93.5% of the *Microcystis* reads in the metagenome samples. Additionally, we observed that the total coverage using all the *Microcystis* genotypes (alignment identity cutoff = 99%) and the total coverage using a single *Microcystis* genome as a reference (alignment identity cutoff = 96%) are nearly perfectly correlated (correlation coefficient *R*^*2*^ = 1, *P* < 2.2e-16) (Spearman correlation) (Supplementary Fig. 13).

### *Microcystis* genotypic diversity variation according to environmental variables

To determine the variables that explain the variation in *Microcystis* community composition, we used a dataset of 42 metagenomes and 14 genotypes from Lake Champlain. Metagenomes with incomplete metadata were excluded. We focused on Lake Champlain as we observed a greater diversity of *Microcystis* genotypes compared to Brazil, including both microcystin-producing and non-producing genotypes. We first used a distance-based redundancy analysis (dbRDA) with the square root of the Bray Curtis distance matrix to investigate *Microcystis*–environment relationships^69,70^ (capscale function from vegan R package, (v2.5.6l)^71^). Variables were pre-selected using the ordiR2step R function^72^ (See Supplementary Methods). The environmental matrix variables included: total phosphorus in μg/l (TP), total nitrogen in μg/l (TN), soluble reactive phosphorus in μg/l (DP), dissolved nitrogen in μg/l (DN), 1-week-cumulative precipitation in mm, 1-week-average air temperature in Celsius, temporal variables (Years, Months and Season) and sampling sites within Lake Champlain (Pelagic or Littoral) (Supplementary Table 3)^43^. To determine the significance of constraints, we used the anova.cca() function from the R vegan package.

We also calculated the beta diversity between groups of samples using the Phyloseq R package (v1.30.0) and the square root of Bray Curtis distance. We used nonmetric multi-dimensional scaling (NMDS, from the phyloseq package that incorporates the metaMDS() function from the R vegan^71,73,74^ package to ordinate the data. Differences in community structure between groups were tested using permutational multivariate analysis of variance (PERMANOVA^75^) with the adonis() function. As PERMANOVA tests might be sensitive to dispersion, we also tested for dispersion by performing an analysis of multivariate homogeneity (PERMDISP^76^) with the permuted betadisper() function.

### Identifying the correlation between microbiome members and *Microcystis* in freshwater samples from Canada

Using the 59 species identified in the *Microcystis* microbiome from Canada and the software MIDAS (v1.3.0), we built a custom gene marker database of 15 universal single-copy gene families. This database also included a reference genome from *Microcystis* (M083S1) and two *Dolichospermum* reference genomes (*D. circinale* AWQC131C and AWQC310F). Using MIDAS, we estimated the relative abundances, reads count, and the read coverage of each associated bacterial species in 72 shotgun metagenomes from Quebec, Canada. Reads were mapped against the custom database including the associated bacteria species. A cuff-off value of nucleotide identity greater or equal to 96% was used for the read mapping. By merging the values (coverage and read counts) for species within the same genus, obtained coverage and read counts at the genus level, for 32 genera of associated bacteria. We used the Spearman rank-based correlation to investigate patterns of co-occurrence between *Microcystis*, *Dolichospermum* and the associated bacterial species and genera in environmental metagenomes. First, the read counts in the matrices containing the genera and species were used to estimate the correlation values (*r*) and p-values between pair of species or genera by using the rcorr() function of the Hmisc (v4.3.0) R package^77^. We also calculated Spearman correlations on the coverage values, yielding similar results. *P-* values were corrected to control the false discovery rate using the qvalue() function from the qvalue (v2.18.0) R package. We also estimated the correlation between *Microcystis* and the AB using the software FastSpar (v0.0.10)^78^. This method is a faster implementation of the Sparse Correlation for Compositional Data algorithm (SparCC)^79^. The significance of the test was evaluated using 100 permutations and a bootstrap of 1000. In general, the most prevalent AB taxa in *Microcystis* colonies had significant correlation (*P* < 0.05) with *Microcystis* using both Spearman and SparCC.

### Co-phylogeny between *Microcystis* and the associated microbiome

The nine most prevalent associated bacterial genera were selected for co-phylogeny analysis, which would be underpowered to detect phylogenetic associations with low-prevalence bacteria (*i.e.* small phylogenies). Core genomes were generated using panX and core alignments were computed as described above, for each associated bacterial genus. Phylogenic core genome trees were built individually for each genus using RAxML^61^. Patristic distances (pairwise distances between pairs of tips on a tree) for the *Microcystis* and associated bacteria phylogenies were estimated using the cophenetic.phylo() function from the ape R-package^68^. The *Microcystis* core genome tree and the tree of the associated bacteria were compared using Parafit test (parafit() function of the ape R package) (See Supplementary Methods)^68,80^. Co-phylogeny trees were built using the function cophylo() from the phytools R package^81^.

### Recent HGT between *Microcystis* and associated bacteria (AB)

To infer recent horizontal gene transfer (HGT) events between *Microcystis* and associated bacteria, we first inferred the pangenomes for each combination of one AB and *Microcystis*, and repeated this for the 72 associated bacterial species. Core and accessory genes with a minimum percentage identity for blastp equal to 99% were identified. We retained those clusters of genes present in at least four genomes, and present in both AB and *Microcystis*. The remaining putatively horizontal transferred genes were annotated in 23 COG (clusters of orthologous groups) categories using eggNOG-mapper (v2.0.1)^82^. Using the package STAMP (v2.1.3) and a chi-square test, we estimated if there were statistical differences in the COG categories between *Microcystis* core genes and the putative horizontally transferred genes^83^. P-values were corrected using Benjamini-Hochberg (controlling the false discovery rate) method. We also estimated HGT events between *Microcystis* and associated species using a second method, Metachip (v1.8.2) (default parameters). The Metachip approach uses both the best match approach (blastn) and a phylogenetic approach to infer HGT (reconciliation between a gene tree and its species tree)^37^.

### Gene functional annotation

The *Microcystis* and associated bacteria genomes were functionally annotated using enrichM (v0.5.0) (https://github.com/geronimp/enrichM)^84^. A PCA based on the presence/absence of KEGG Orthologous genes (KO) in *Microcystis* and associated bacteria genera was generated using the option ‘enrichment’ in enrichM. Genome groups (*Microcystis* vs each associated bacteria genus) were compared using the same option. KEGG modules differentially abundant in *Microcystis* or the associated bacteria genus were filtered based on a completeness greater or equal to 70%.

*Microcystis* and associated bacterial genomes (109 *Microcystis* and 391 associated genomes) were annotated using Roary (v3.13.0). The resulting genomes in GenBank format were used to predict the biosynthetic gene clusters (BGCs) using default parameters (--taxon bacteria --cb-general --cb-knownclusters --cb-subclusters --asf --pfam2go --smcog-trees --genefinding-tool prodigal-m) in antiSMASH (v5.1.2)^85,86^. The BIG-SCAPE package (v1.0.1) with default parameters analysed the ANTISMASH BGCs and based on a similarity network classified them into Gene Cluster Families (GCFs)^87^. BGCs were classified in BiG-SCAPE classes (*e.g.*, polyketide synthases nonribosomal peptide synthetases (NRPSs), post-translationally modified peptides (RiPPs) and terpenes. A total of 2,395 BGCs were identified in 415 genomes.

## Supporting information

R script 1

R script 2

## Data availability

Raw sequences and metagenome assembled genomes (MAGs) are available in NCBI under Bioproject numbers PRJNA507251 and PRJNA662092.

## Acknowledgements

We are grateful to Julie Marleau, Miria Elias and Alberto Mazza for assistance in the sampling. This work was supported by the Genome Québec and Genome Canada-funded ATRAPP Project (Algal blooms, Treatment, Risk Assessment, Prediction and Prevention). Colonies and water samples from Brazil were obtained thanks to a FAPEMIG grant to A.G. We also want to acknowledge the financial support of the National Research Council.

## Author contributions

B.J.S., N.T. and O.M.P.C. designed the study. O.M.P.C., N.T., A.G., L.C.B.M. and N.F. performed the lab experiments. N.T. and O.M.P.C. performed the data analyses. E.M. and O.M.P.C. performed the cophylogeny. B.J.S., N.T. and O.M.P.C. wrote the manuscript. B.J.S., N.T., O.M.P.C., A.G., Y.T. and N.F. contributed to its reviewing and editing.

