## Supplementary Figures for "Single-colony sequencing reveals phylosymbiosis, co-phylogeny, and horizontal gene transfer between the cyanobacterium *Microcystis* and its microbiome"

**Supplementary figure legends**

**Figure 1.** ML Phylogenetic tree of *Microcystis* genomes (a) and ANIm clusters (b) corresponding to monophyletic clades with ANI values greater or equal to 99%.

**Figure 2.** Phylogenetic tree of *Microcystis* and 72 associated species (391 MAGs) recovered from *Microcystis* colonies. Taxonomic annotation for the genomes to different genera and species was done using Blastp of the RecA and RpoB proteins against the NCBI database and then complemented using the Genome Taxonomy Database Toolkit (GTDB-Tk) v1.0.2. The colored circles indicate the most prevalent genera in the *Microcystis* colony Microbiome.

**Figure 3.** Associated bacteria prevalence in the *Microcystis* colony microbiome from Canada and Brazil. Based on the normalized read coverages greater or equal than 1%, we recalculated the richness or number of associated species to the *Microcystis* colonies.

**Figure 4.** *Microcystis* genotype diversity over time in environmental metagenomes from Lake Champlain. Using MIDAS software, we estimated the relative abundances (a) and the read coverage (b) of the *Microcystis* genotypes in 72 shotgun metagenomes from Lake Champlain, Quebec (62 metagenomes from a long-term experiment (2006 to 2016), plus 10 metagenomes from 2017 and 2018). The metagenomes with *Microcystis* genotype coverage lower than 1 or incomplete metadata were excluded.

**Figure 5.** *Microcystis* genotypic diversity variation over time. We used dbRDA (a) and NMDS (b) to ordinate the metagenome samples from Quebec based on the Bray–Curtis dissimilarities of the normalized coverage of *Microcystis* genotypes.

**Figure 6.** *Microcystis* genotype diversity over time in environmental metagenomes from Pampulha reservoir. The genotypes containing the mcy cluster are indicated with an asterisk.

**Figure 7.** Associated bacteria (AB) diversity variation over time. We used dbRDA (a) and non- NMDS (b) to ordinate the metagenome samples from Quebec based on the Bray–Curtis dissimilarities of the normalized coverage of the AB species.

**Figure 8.** *Microcystis*, *Dolichospermum* and associated bacteria species coverage in environmental metagenomes across time. Figure 8a shows the *Microcystis* and *Dolichospermum* normalized read coverages across time. Figure 8b shows the *Microcystis*-associated bacteria genus read coverage.

**Figure 9.** Certain associated bacteria species are well correlated with the presence of *Microcystis*. Spearman correlation coefficients were estimated between *Microcystis* and each associated bacterium and between *Dolichospermum* and each associated bacterium. Figure 9a shows the comparisons at genus level and Figure 9b the comparisons at species level. The axis x was sorted according with the genus and species prevalence in the *Microcystis* colonies from Canada. The correlations between the AB and *Microcystis* were also estimated using fastspar (Figure 9c and 9d).

**Figure 10**. *Microcystis* genotype-specific associations with associated bacteria species in environmental metagenomes. Normalized read counts were estimated per each *Microcystis* genotype and the associated bacteria species in 72 environmental metagenomes. The Spearman correlation coefficients were estimated between each *Microcystis* genotype and each associated bacteria species (Figure 10a). The correlations were also estimated using fastspar (Figure 10b).

**Figure 11.** PCA based on the presence absence of KEGG Orthologous genes in *Microcystis* and associated bacteria genera.

**Figure 12.** MIDAS workflow used to analyze the *Microcystis* colony and metagenome community composition.

**Figure 13.** Pearson correlation between *Microcystis* genotype coverage with alignment identity of ≥99% and *Microcystis* reference coverage with alignment identity of ≥96%.

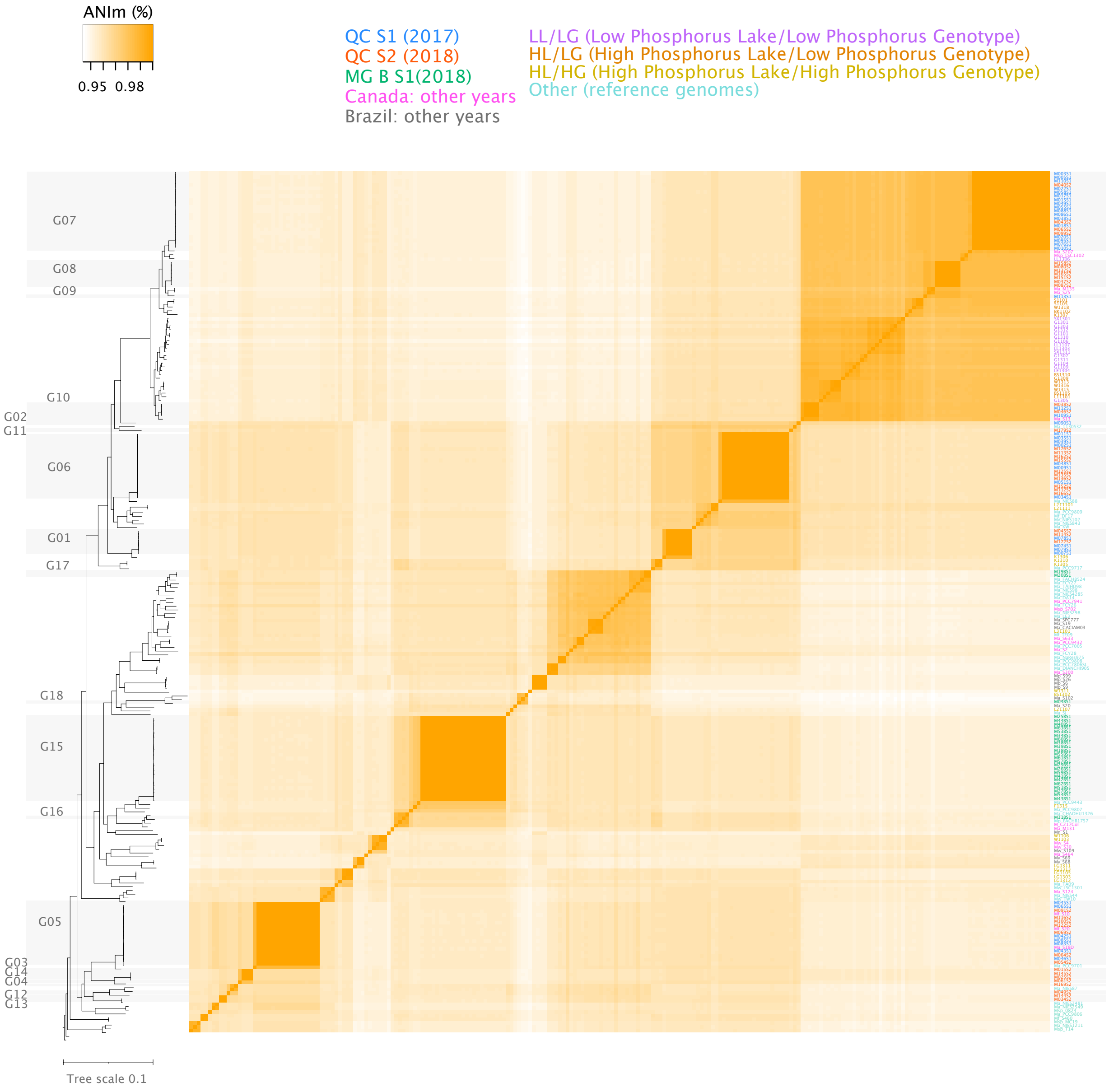
ANIm (%)
0.95 0.98
QC S1 (2017)
QC S2 (2018)
MG B S1(2018)
Canada: other years
Brazil: other years
LL/LG (Low Phosphorus Lake/Low Phosphorus Genotype)
HL/LG (High Phosphorus Lake/Low Phosphorus Genotype)
HL/HG (High Phosphorus Lake/High Phosphorus Genotype)
Other (reference genomes)
G07
G08
G09
G10
G02
G11
G06
G01
G17
G18
G15
G16
G05
G03
G14
G04
G12
G13
Tree scale 0.1

**Fig. S1**

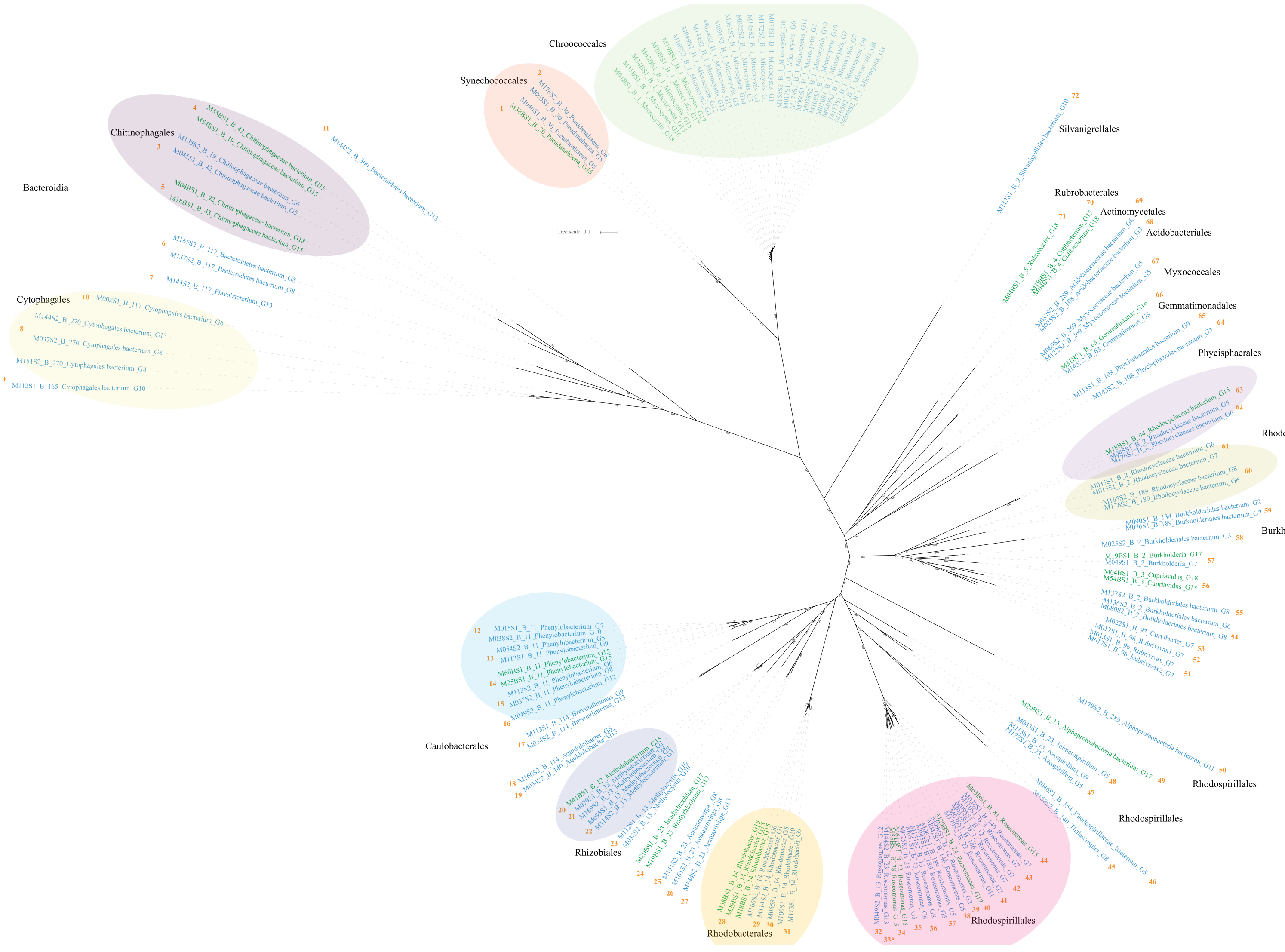
Chroococcales
Synechococcales
Chitinophagales
Bacteroidia
Cytophagales
Silvanigrellales
Rubrobacterales
Actinomycetales
Acidobacteriales
Myxococcales
Gemmatimonadales
Phycisphaerales
Rhodocyclales
Burkholderiales
Caulobacterales
Rhizobiales
Rhodobacterales
Rhodospirillales
Rhodospirillales
Rhodospirillales
Tree scale: 0.1

**Fig. S2**

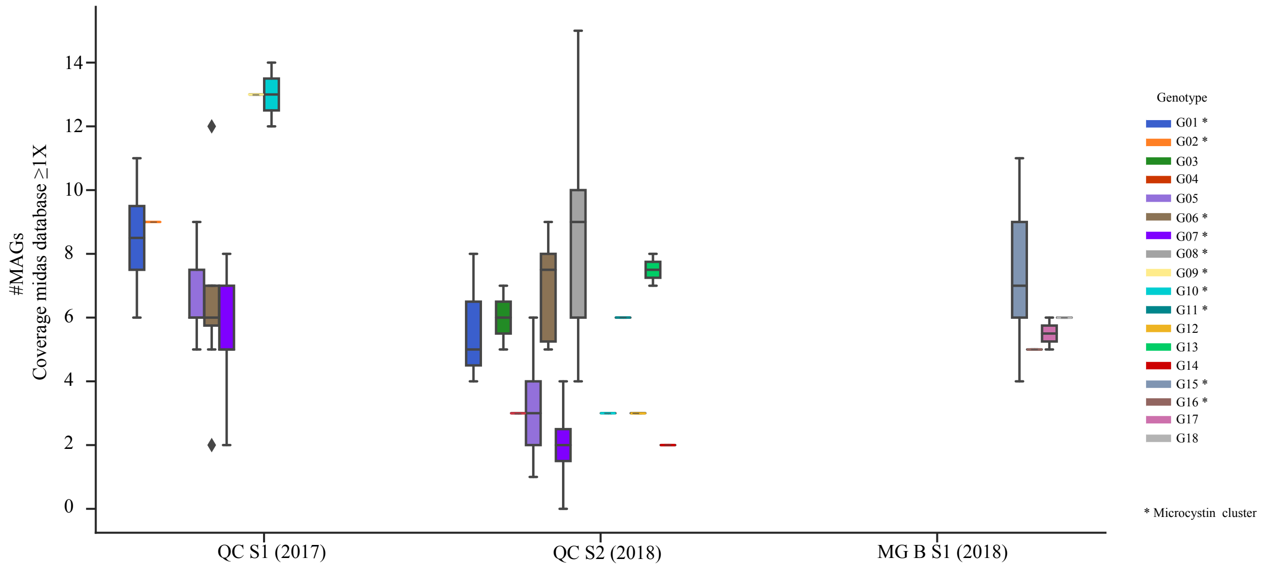

#MAGs
Coverage midas database ≥1X
14
12
10
8
6
4
2
0
QC S1 (2017)
QC S2 (2018)
MG B S1 (2018)
Genotype
G01 *
G02 *
G03
G04
G05
G06 *
G07 *
G08 *
G09 *
G10 *
G11 *
G12
G13
G14
G15 *
G16 *
G17
G18
* Microcystin cluster


**Fig. S3**

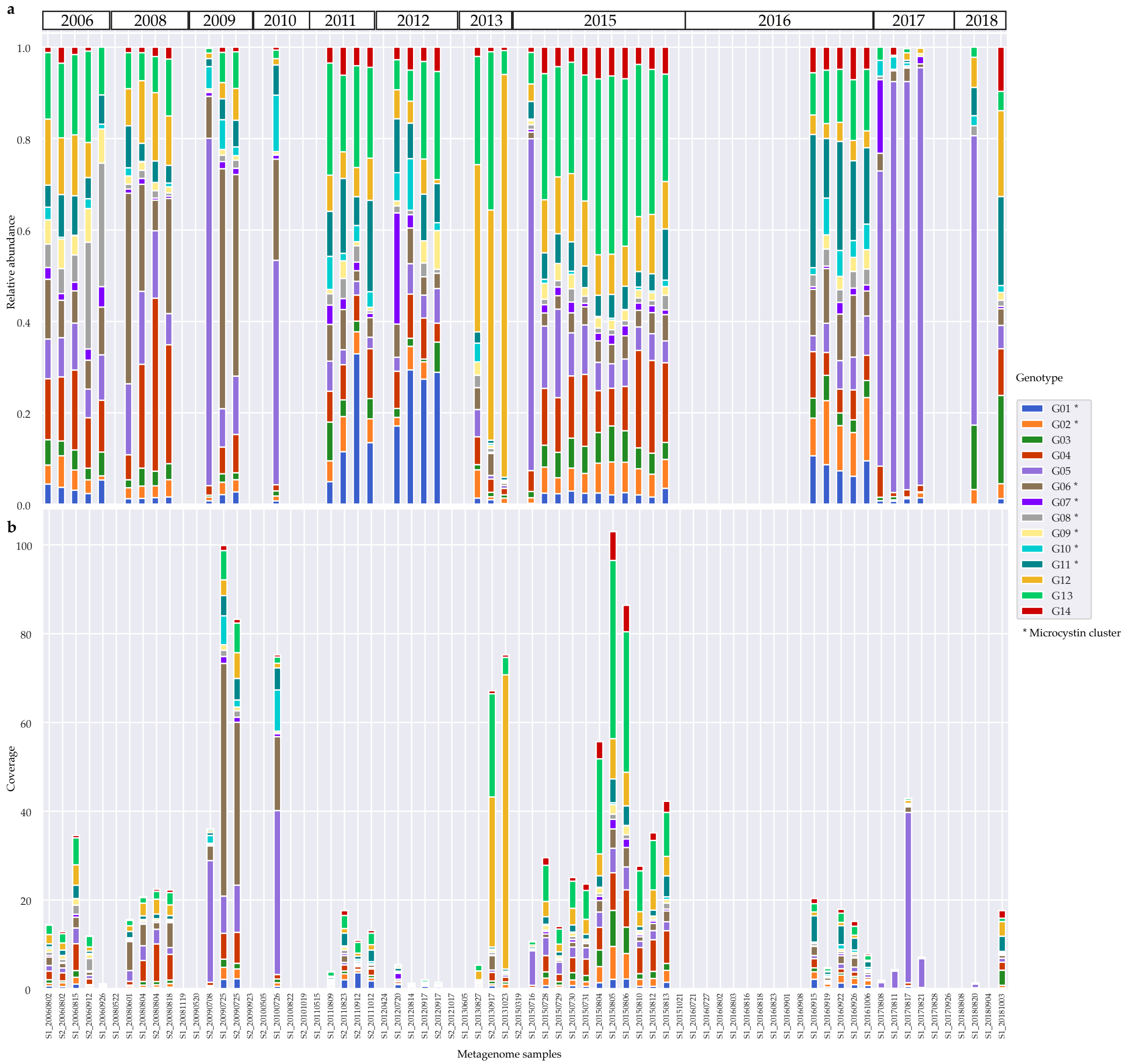
a
b
2006
2008
2009
2010
2011
2012
2013
2015
2016
2017
2018
Relative abundance
Coverage
Metagenome samples
Genotype
G01 *
G02 *
G03
G04
G05
G06 *
G07 *
G08 *
G09 *
G10 *
G11 *
G12
G13
G14
* Microcystin cluster

**Fig. S4**

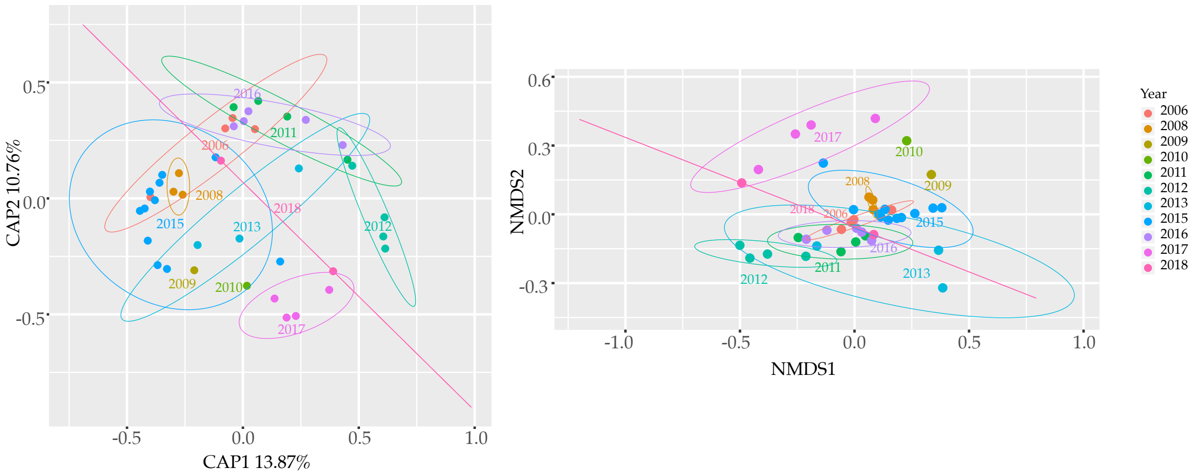
CAP2 10.76%
CAP1 13.87%
NMDS2
NMDS1
Year
2006
2008
2009
2010
2011
2012
2013
2015
2016
2017
2018

**Fig. S5**

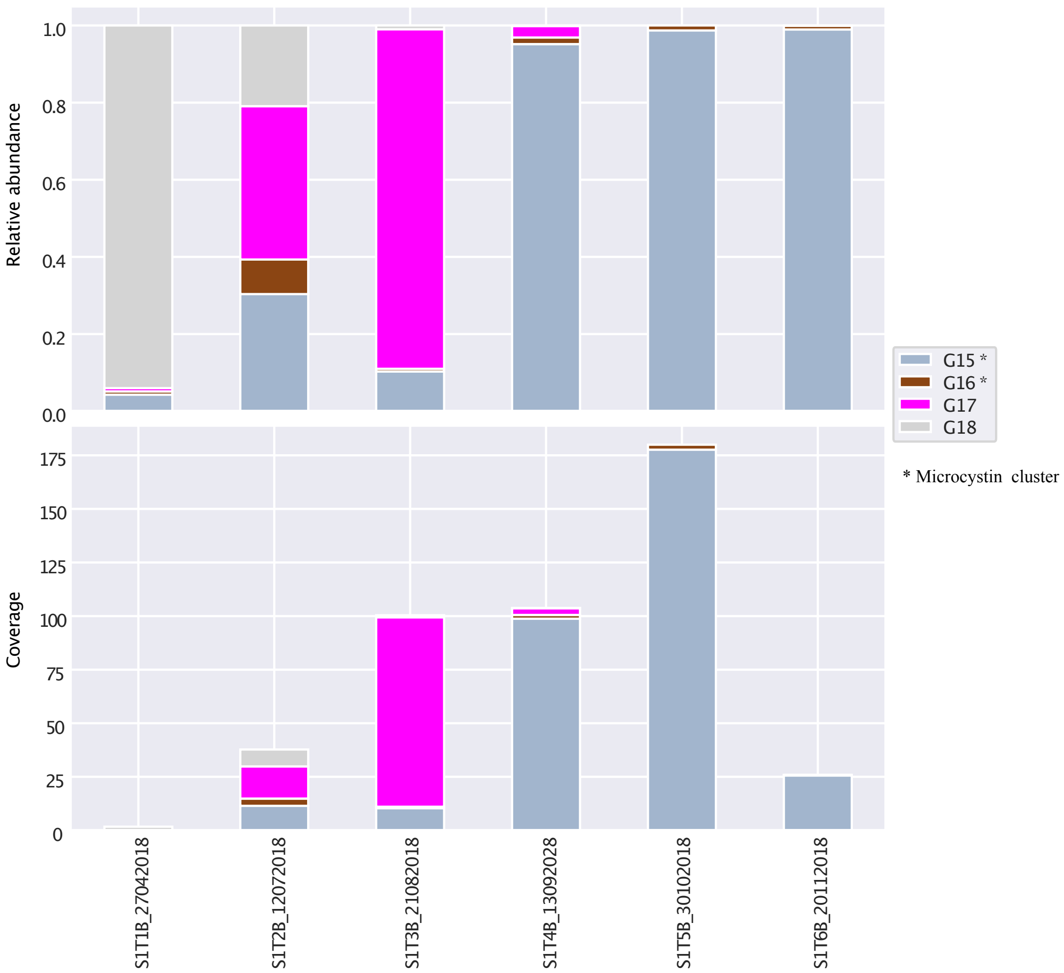
Relative abundance
Coverage
G15 *
G16 *
G17
G18
* Microcystin cluster
S1T1B_27042018
S1T2B_12072018
S1T3B_21082018
S1T4B_13092028
S1T5B_30102018
S1T6B_20112018

**Fig. S6**

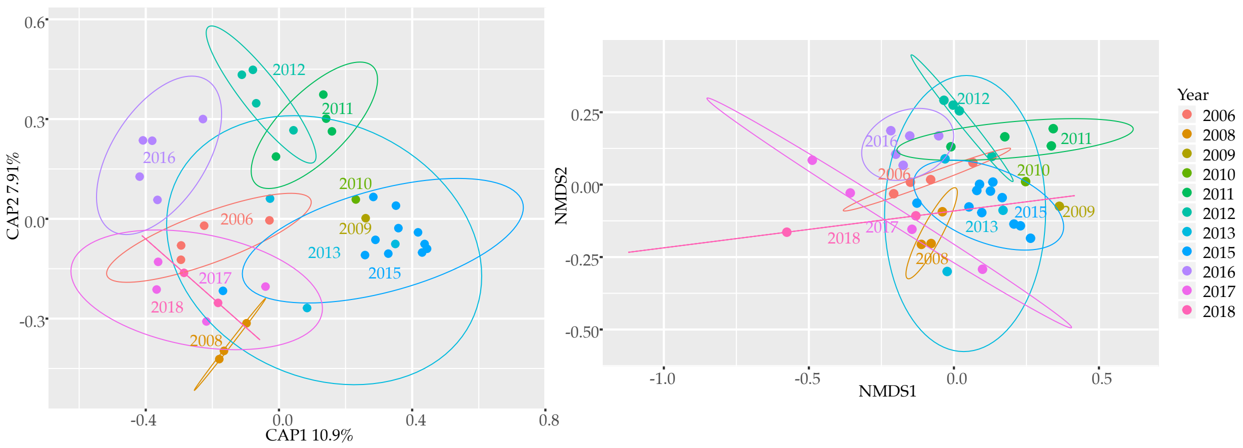
CAP1 10.9%
CAP2 7.91%
NMDS1
NMDS2
Year
2006
2008
2009
2010
2011
2012
2013
2015
2016
2017
2018

### Fig. S7

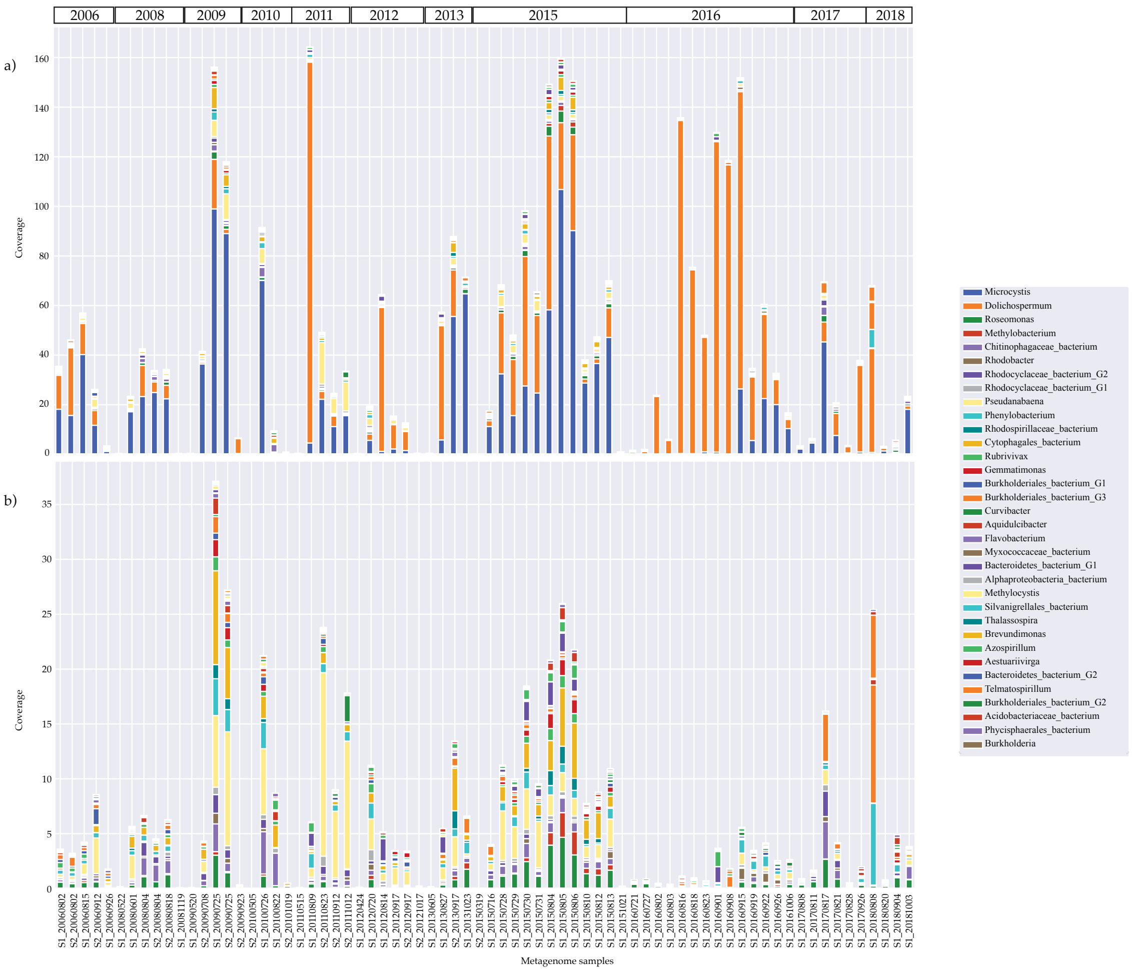
2006
2008
2009
2010
2011
2012
2013
2015
2016
2017
2018
a)
b)
Coverage
Metagenome samples
Microcystis
Dolichospermum
Roseomonas
Methylobacterium
Chitinophagaceae_bacterium
Rhodobacter
Rhodocyclaceae_bacterium_G2
Rhodocyclaceae_bacterium_G1
Pseudanabaena
Phenylobacterium
Rhodospirillaceae_bacterium
Cytophagales_bacterium
Rubrivivax
Gemmatimonas
Burkholderiales_bacterium_G1
Burkholderiales_bacterium_G3
Curvibacter
Aquidulcibacter
Flavobacterium
Myxococcaceae_bacterium
Bacteroidetes_bacterium_G1
Alphaproteobacteria_bacterium
Methylocystis
Silvanigrellales_bacterium
Thalassospira
Brevundimonas
Azospirillum
Aestuariivirga
Bacteroidetes_bacterium_G2
Telmatospirillum
Burkholderiales_bacterium_G2
Acidobacteriaceae_bacterium
Phycisphaerales_bacterium
Burkholderia

### Fig. S8

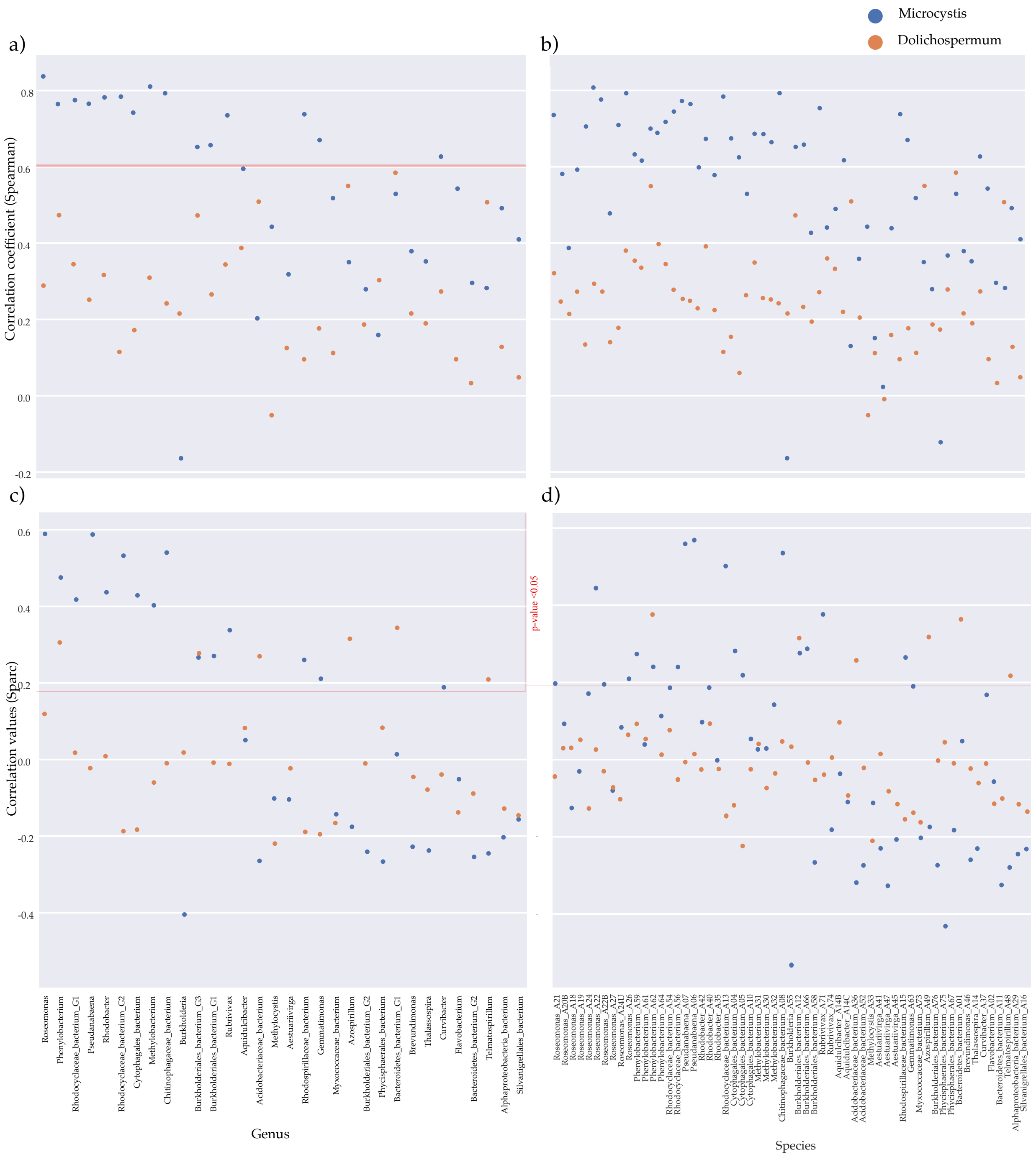

Microcystis
Dolichospermum
a)
b)
c)
d)
Correlation coefficient (Spearman)
Correlation values (Sparc)
p-value <0.05
0.8
0.6
0.4
0.2
0.0
-0.2
-0.4
Roseomonas
Phenylobacterium
Rhodocyclaceae_bacterium_G1
Pseudanabaena
Rhodobacter
Rhodocyclaceae_bacterium_G2
Cytophagales_bacterium
Methylobacterium
Chitinophagaceae_bacterium
Burkholderia
Burkholderiales_bacterium_G3
Burkholderiales_bacterium_G1
Rubrivivax
Aquidulcibacter
Acidobacteriaceae_bacterium
Methylocystis
Aestuariivirga
Rhodospirillaceae_bacterium
Gemmatimonas
Myxococcaceae_bacterium
Azospirillum
Burkholderiales_bacterium_G2
Phycisphaerales_bacterium
Bacteroidetes_bacterium_G1
Brevundimonas
Thalassospira
Curvibacter
Flavobacterium
Bacteroidetes_bacterium_G2
Telmatospirillum
Alphaproteobacteria_bacterium
Silvanigrellales_bacterium
Genus
Roseomonas_A21
Roseomonas_A20B
Roseomonas_A18
Roseomonas_A19
Roseomonas_A24
Roseomonas_A22
Roseomonas_A22B
Roseomonas_A27
Roseomonas_A24U
Roseomonas_A26
Phenylobacterium_A59
Phenylobacterium_A61
Phenylobacterium_A62
Phenylobacterium_A64
Rhodocyclaceae_bacterium_A54
Rhodocyclaceae_bacterium_A56
Pseudanabaena_A07
Pseudanabaena_A06
Rhodobacter_A42
Rhodobacter_A40
Rhodobacter_A35
Rhodocyclaceae_bacterium_A13
Cytophagales_bacterium_A04
Cytophagales_bacterium_A05
Cytophagales_bacterium_A10
Methylobacterium_A31
Methylobacterium_A30
Methylobacterium_A32
Chitinophagaceae_bacterium_A08
Burkholderia_A55
Burkholderiales_bacterium_A12
Burkholderiales_bacterium_A66
Burkholderiales_bacterium_A58
Rubrivivax_A71
Rubrivivax_A74
Aquidulcibacter_A14B
Aquidulcibacter_A14C
Acidobacteriaceae_bacterium_A36
Acidobacteriaceae_bacterium_A52
Methylocystis_A33
Aestuariivirga_A41
Aestuariivirga_A47
Aestuariivirga_A45
Rhodospirillaceae_bacterium_A15
Gemmatimonas_A63
Myxococcaceae_bacterium_A73
Azospirillum_A49
Burkholderiales_bacterium_A76
Phycisphaerales_bacterium_A75
Phycisphaerales_bacterium_A67
Bacteroidetes_bacterium_A01
Brevundimonas_A46
Thalassospira_A14
Curvibacter_A37
Flavobacterium_A02
Bacteroidetes_bacterium_A11
Telmatospirillum_A48
Alphaproteobacteria_bacterium_A29
Silvanigrellales_bacterium_A16
Species


**Fig. S9**

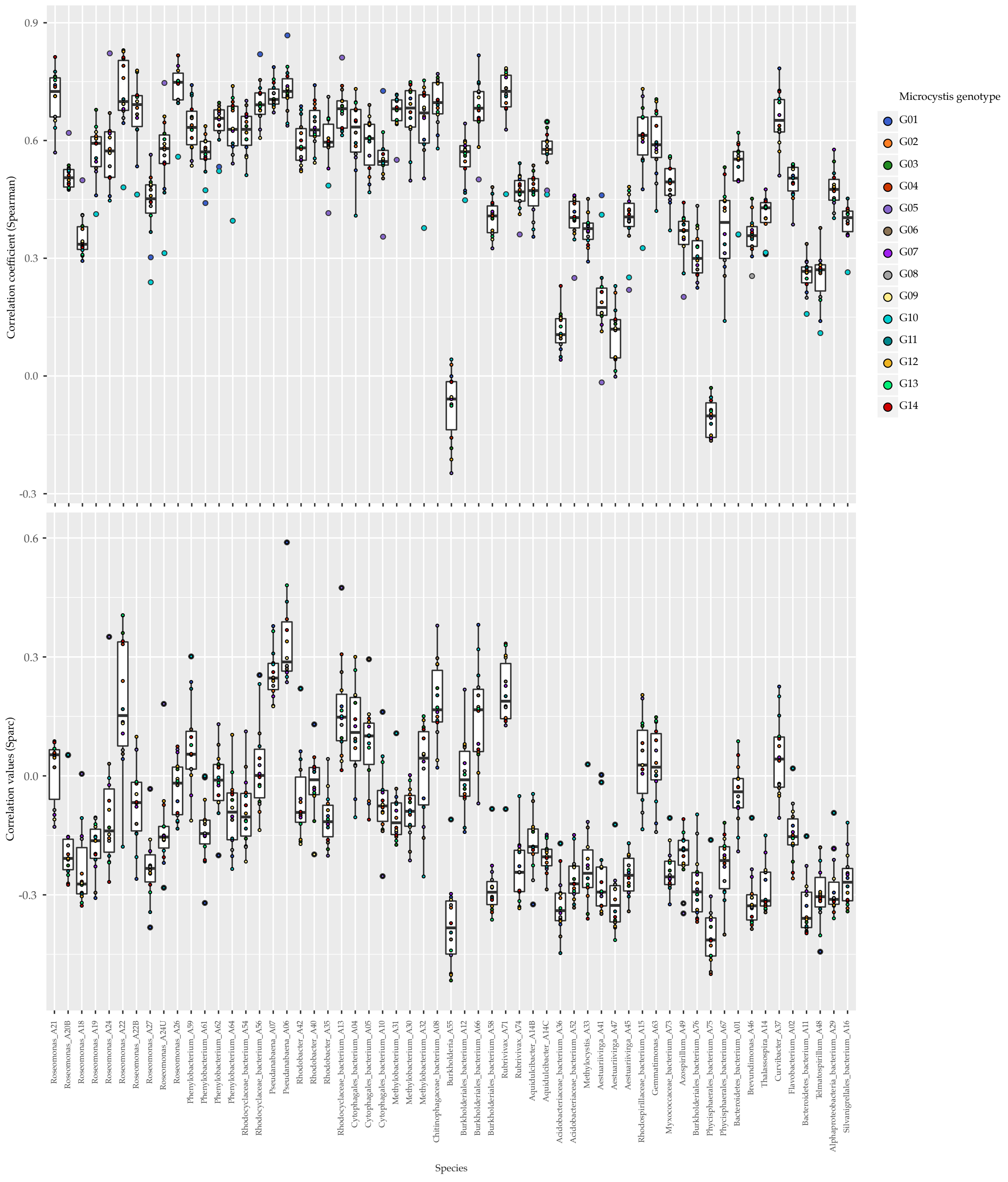

Correlation coefficient (Spearman)
0.9
0.6
0.3
0.0
-0.3
Correlation values (Sparc)
0.6
0.3
0.0
-0.3
Microcystis genotype
G01
G02
G03
G04
G05
G06
G07
G08
G09
G10
G11
G12
G13
G14
Roseomonas_A21
Roseomonas_A20B
Roseomonas_A18
Roseomonas_A19
Roseomonas_A24
Roseomonas_A22
Roseomonas_A22B
Roseomonas_A27
Roseomonas_A24U
Roseomonas_A26
Phenylobacterium_A59
Phenylobacterium_A61
Phenylobacterium_A62
Phenylobacterium_A64
Rhodocyclaceae_bacterium_A54
Rhodocyclaceae_bacterium_A56
Pseudanabaena_A07
Pseudanabaena_A06
Rhodobacter_A42
Rhodobacter_A40
Rhodobacter_A35
Rhodocyclaceae_bacterium_A13
Cytophagales_bacterium_A04
Cytophagales_bacterium_A05
Cytophagales_bacterium_A10
Methylobacterium_A31
Methylobacterium_A30
Methylobacterium_A32
Chitinophagaceae_bacterium_A08
Burkholderia_A55
Burkholderiales_bacterium_A12
Burkholderiales_bacterium_A66
Burkholderiales_bacterium_A58
Rubrivivax_A71
Rubrivivax_A74
Aquidulcibacter_A14B
Aquidulcibacter_A14C
Acidobacteriaceae_bacterium_A36
Acidobacteriaceae_bacterium_A52
Methylocystis_A33
Aestuariivirga_A41
Aestuariivirga_A47
Aestuariivirga_A45
Rhodospirillaceae_bacterium_A15
Gemmatimonas_A63
Myxococcaceae_bacterium_A73
Azospirillum_A49
Burkholderiales_bacterium_A76
Phycisphaerales_bacterium_A75
Phycisphaerales_bacterium_A67
Bacteroidetes_bacterium_A01
Brevundimonas_A46
Thalassospira_A14
Curvibacter_A37
Flavobacterium_A02
Bacteroidetes_bacterium_A11
Telmatospirillum_A48
Alphaproteobacteria_bacterium_A29
Silvanigrellales_bacterium_A16
Species


**Fig. S10**

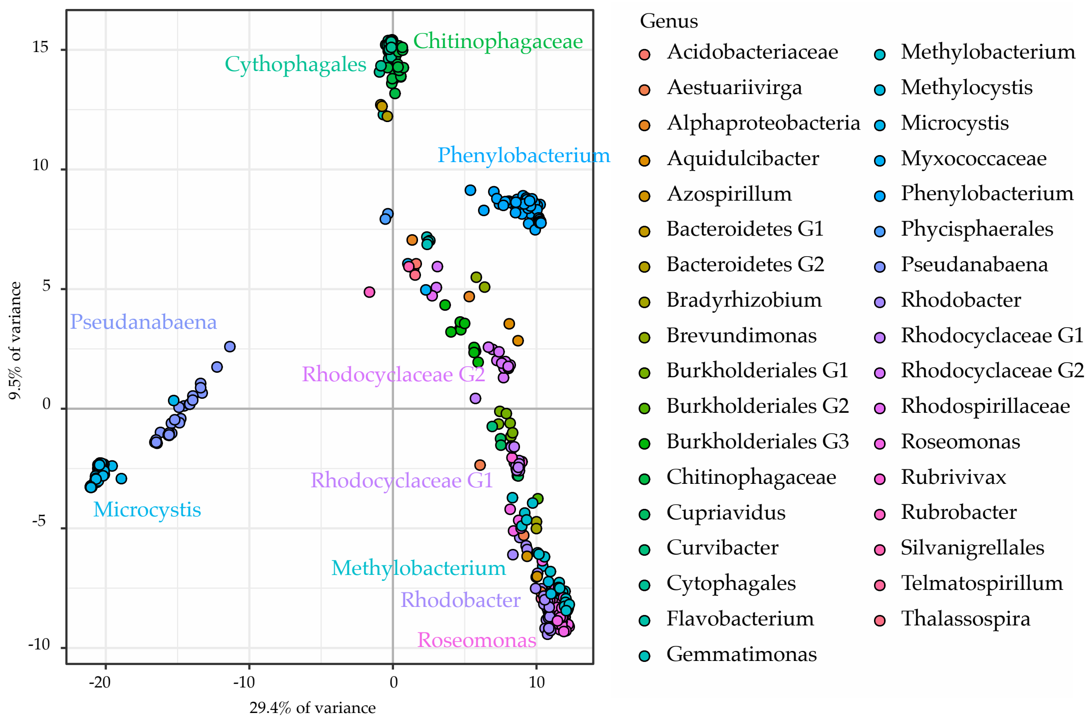
Chitinophagaceae
Cythophagales
Phenylobacterium
Pseudanabaena
Rhodocyclaceae G2
Rhodocyclaceae G1
Microcystis
Methylobacterium
Rhodobacter
Roseomonas
9.5% of variance
29.4% of variance
Genus
Acidobacteriaceae
Aestuariivirga
Alphaproteobacteria
Aquidulcibacter
Azospirillum
Bacteroidetes G1
Bacteroidetes G2
Bradyrhizobium
Brevundimonas
Burkholderiales G1
Burkholderiales G2
Burkholderiales G3
Chitinophagaceae
Cupriavidus
Curvibacter
Cytophagales
Flavobacterium
Gemmatimonas
Methylobacterium
Methylocystis
Microcystis
Myxococcaceae
Phenylobacterium
Phycisphaerales
Pseudanabaena
Rhodobacter
Rhodocyclaceae G1
Rhodocyclaceae G2
Rhodospirillaceae
Roseomonas
Rubrivivax
Rubrobacter
Silvanigrellales
Telmatospirillum
Thalassospira

**Fig. S11**

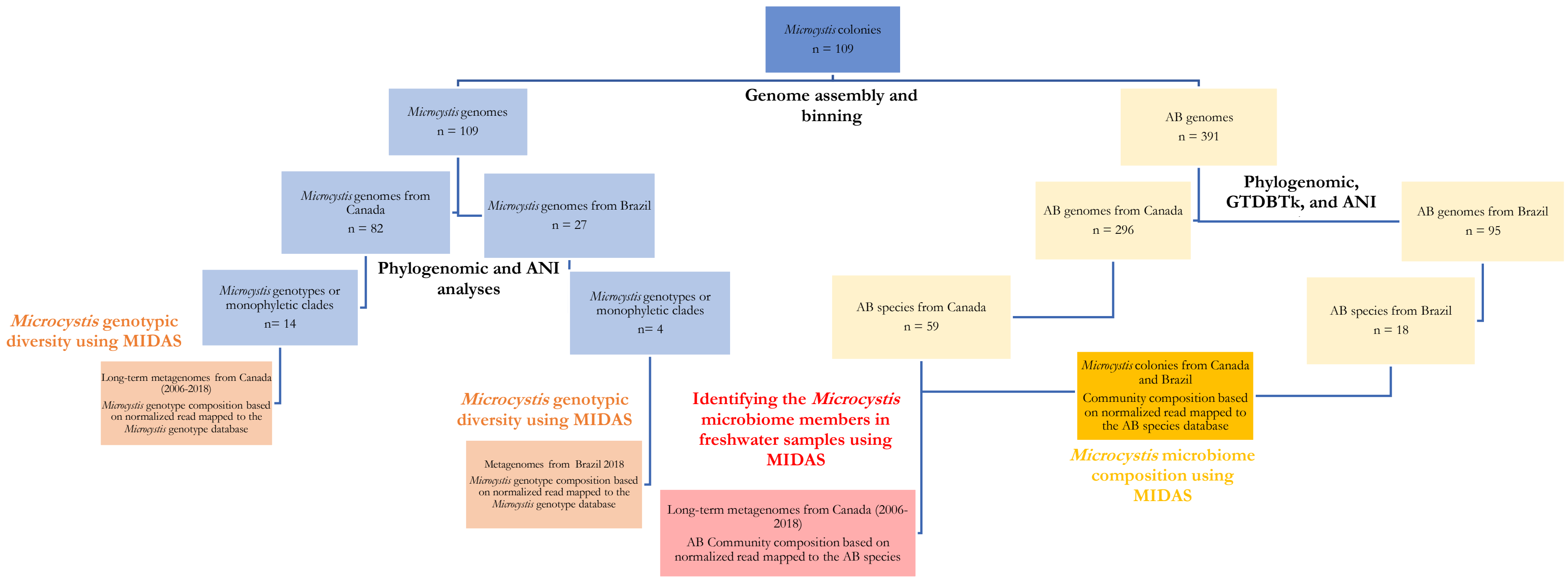

Microcystis colonies
n = 109
Genome assembly and binning
Microcystis genomes
n = 109
AB genomes
n = 391
Microcystis genomes from Canada
n = 82
Microcystis genomes from Brazil
n = 27
Phylogenomic, GTDBTk, and ANI
AB genomes from Canada
n = 296
AB genomes from Brazil
n = 95
Phylogenomic and ANI analyses
Microcystis genotypes or monophyletic clades
n= 14
Microcystis genotypes or monophyletic clades
n= 4
AB species from Canada
n = 59
AB species from Brazil
n = 18
Microcystis genotypic diversity using MIDAS
Long-term metagenomes from Canada (2006-2018)
Microcystis genotype composition based on normalized read mapped to the Microcystis genotype database
Microcystis genotypic diversity using MIDAS
Metagenomes from Brazil 2018
Microcystis genotype composition based on normalized read mapped to the Microcystis genotype database
Identifying the Microcystis microbiome members in freshwater samples using MIDAS
Long-term metagenomes from Canada (2006-2018)
AB Community composition based on normalized read mapped to the AB species
Microcystis colonies from Canada and Brazil
Community composition based on normalized read mapped to the AB species database
Microcystis microbiome composition using MIDAS


**Fig. S12**

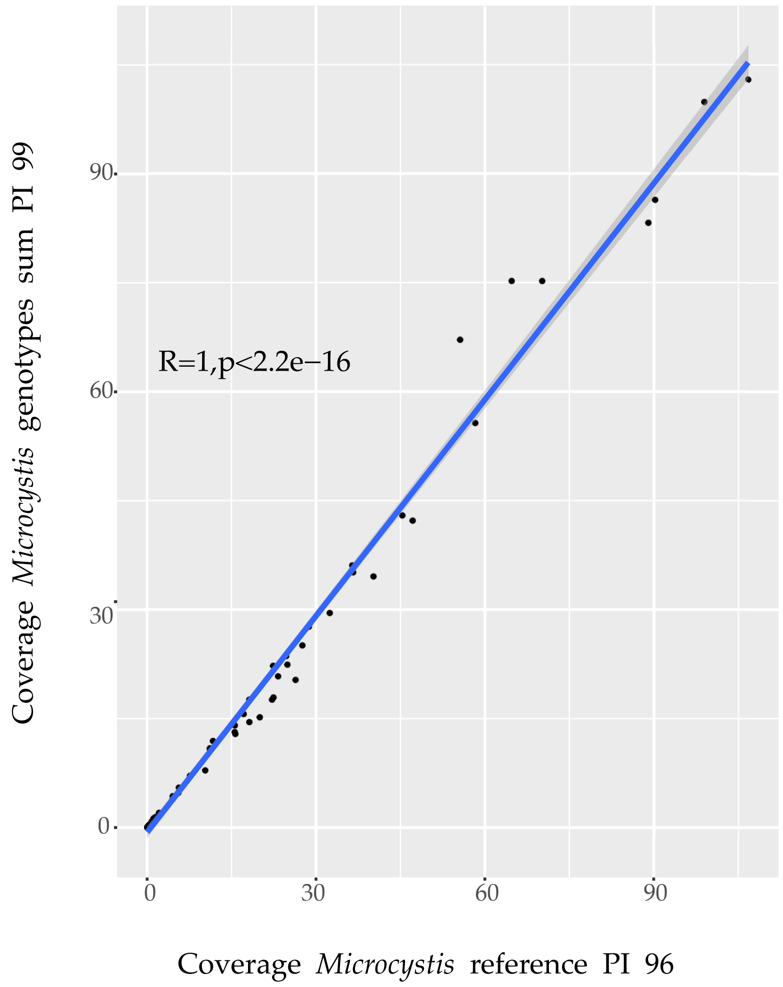

R=1,p<2.2e−16
Coverage *Microcystis* genotypes sum PI 99
0
30
60
90
Coverage *Microcystis* reference PI 96


**Fig. S13**
