## Supplementary Methods for "Single-colony sequencing reveals phylosymbiosis, co-phylogeny, and horizontal gene transfer between the cyanobacterium *Microcystis* and its microbiome"

1. **Microcystis colonies isolation**

**Isolation of the Microcystis colony using micropipes (1 to 1000 ul), sterile medium (Z8 medium) and a microscope.**

We washed the colonies 15-20 times using sterile Z8 medium^1^ and stored them in 20 ul of Z8 liquid medium in Eppendorf tubes (1.5 ml) at -80 C until the DNA extraction.

Colonies sizes were greater than 0.6 mm. We also found big colonies with size greater than 3 mm.

2. **Microcystis colonies DNA isolation**

**We used the ChargeSwitch® gDNA Mini Bacteria Kit (Catalog number: CS11301 -** **Invitrogen -Thermo Fisher Scientific) for the isolating genomic DNA**

**The original protocol for DNA isolation was modified in the following steps.**

2.2 **Bacterial Cell Lysis protocol**

**We skipped the steps 1 and 2.**

Before starting at the step 3. Microcystis colony tubes were kept in ice. Then Microcystis colony were slowly thaw and added to a tube containing 50 mg of beads (PowerBead Tubes of 2 mL, Glass bead size 0.1 mm, Cat No./ID: 13118-50 - Qiagen).

**Step 3.**

We started the cell lysis protocol in the step 3 (See pages 12-16 <https://www.thermofisher.com/document-connect/document-connect.html?url=https%3A%2F%2Fassets.thermofisher.com%2FTFS-Assets%2FLSG%2Fmanuals%2Fchargeswitch_minibacteria_man.pdf>). Please noticed that we did not use lysostaphin. The lysozyme used correspond to the following reference: Lysozyme from chicken egg white (Catalog Number L6876 from Sigma-Aldrich®)

**Step 4.** We incubated the sample for 20 minutes instead of 10 minutes at 37°C.

**Step 5.** We kept it as in the user guide.

**Step 6.** We kept it as in the user guide.

**Step 7.** We incubated the sample for 1 hour at 55 °C.

During the incubation, we inverted gently the capped tube at least six times every ten minutes. Please make sure that the colony stays in the solution.

After the incubation, we vortexed the tube using the tissue lyse for 3 minutes (45 oscillation per second or max speed). (TissueLyser LT from Qiagen). Then we centrifuged the tube for 1 minute at 9000 rcf to remove the bubbles.

1.2 **Binding DNA protocol**

**Step 1.** We vortexed the ChargeSwitch® Magnetic Beads for 20 seconds.

**Step 2.** We kept it as in the user guide.

**Step 3.** We kept it as in the user guide.

**Step 4.** We kept it as in the user guide.

**Step 5.** We used two minutes instead of one minute.

During step 5, we transferred the mix to a mini centrifuge tube (1.5 ml). Please make sure of transfer all the volume, including the glass beads. Continue with step 6.

**Step 6**. We used two minutes instead of one minute.

**Step 7.** We kept it as in the user guide.

**Step 8.** We kept it as in the user guide.

1.3 **Washing DNA protocol**

We kept all the steps as in the user guide, except for steps 3 and 4. In step 3 we used two minutes instead of one minute. In the steps 4 we also tried to remove carefully the glass beads at the bottom of the tube.

1.4 **Eluting DNA protocol**

We kept all the steps as in the user guide, except for the steps 2 and 3.

**Step 2.** We added 40 ul of Elution Buffer instead of Add 200 μL.

**Step 3.** We incubated the sample at 65 °C for 15 minutes.

**R script 1**

**R script 2**
