## Supplementary material for "Single-colony sequencing reveals phylosymbiosis, co-phylogeny, and horizontal gene transfer between the cyanobacterium *Microcystis* and its microbiome": Table S10

Table S10. Colonies sequenced per each sampling time point.

|  | Number of sequenced colonies | Time | Region |
| --- | --- | --- | --- |
| Sample 1 | 12 | 08-Aug-2017 | Station 1 or Littoral, Lake Champlain, Quebec, Canada |
| Sample 2 | 4 | 11-Aug-2017 | Station 1 or Littoral, Lake Champlain, Quebec, Canada |
| Sample 3 | 6 | 17-Aug-2017 | Station 1 or Littoral, Lake Champlain, Quebec, Canada |
| Sample 4 | 7 | 21-Aug-2017 | Station 1 or Littoral, Lake Champlain, Quebec, Canada |
| Sample 5 | 7 | 28-Aug-2017 | Station 1 or Littoral, Lake Champlain, Quebec, Canada |
| Sample 6 | 4 | 26-Sep-2017 | Station 1 or Littoral, Lake Champlain, Quebec, Canada |
| Sample 7 | 3 | 08-Aug-2018 | Station 1 or Littoral, Lake Champlain, Quebec, Canada |
| Sample 8 | 13 | 20-Aug-2018 | Station 1 or Littoral, Lake Champlain, Quebec, Canada |
| Sample 9 | 9 | 04-Sep-2018 | Station 1 or Littoral, Lake Champlain, Quebec, Canada |
| Sample 10 | 9 | 20-Sep-2018 | Station 1 or Littoral, Lake Champlain, Quebec, Canada |
| Sample 11 | 8 | 03-Oct-2018 | Station 1 or Littoral, Lake Champlain, Quebec, Canada |
| Sample 12 | 1 | 27-Apr-2018 | Pampulha reservoir, Minas Gerais, Brazil |
| Sample 13 | 3 | 21-Aug-2018 | Pampulha reservoir, Minas Gerais, Brazil |
| Sample 14 | 5 | 13-Sep-2018 | Pampulha reservoir, Minas Gerais, Brazil |
| Sample 15 | 10 | 30-Oct-2018 | Pampulha reservoir, Minas Gerais, Brazil |
| Sample 16 | 8 | 20-Nov-2018 | Pampulha reservoir, Minas Gerais, Brazil |
| Total | 109 |  |  |
